# The emergence and loss of cyclic peptides in *Nicotiana* illuminate dynamics and mechanisms of plant metabolic evolution

**DOI:** 10.1101/2024.11.29.626059

**Authors:** Elliot M Suh, Jakob K Reinhardt, Jing-Ke Weng

## Abstract

Specialized metabolism plays a central role in how plants cope with both biotic and abiotic stresses in order to survive and reproduce within dynamic and challenging environments. One recently circumscribed class of plant-specific, ribosomally synthesized and post-translationally modified peptides are the burpitides, which are characterized by the installation of distinct sidechain macrocycles by enzymes known as burpitide cyclases. While they are found across many plant families and exhibit diverse bioactivities, little is known about their evolution or how new variants arise. Here we present the discovery of a new burpitide cyclase, resurrected from a defunct pseudogene from the model organism *Nicotiana attenuata*, the coyote tobacco. By repairing the pseudogene *ΨNatBURP2* and expressing it heterologously in *Nicotiana benthamiana*, we successfully reconstituted its original enzymatic activity. As an autocatalytic peptide cyclase, it installs a unique C-C bond between the tyrosine side chain and a specific backbone a-carbon of a heptapeptide core motif, resulting in the burpitide dubbed "nanamin." Despite its pseudogenization in *N. attenuata*, we found that the closely related species, *N. clevelandii*, retains the wild-type gene and produces nanamins. Phylogenetic analyses and targeted mutagenesis experiments reveal that this chemotype must have evolved from the duplication and neofunctionalization of a more promiscuous ancestral gene. This work highlights how novel peptide chemotypes may rapidly emerge and disappear in plants, while expanding the molecular toolkit for engineering novel peptides with applications in agriculture and drug discovery.

**Significance:** While RiPPs represent a major source of antibiotics and bioactive compounds, much research has focused on microbial sources even as plant RiPPs go understudied. Here, we resurrect an extinct peptide cyclase from the coyote tobacco through analysis of its functional relatives in other species. This newly identified cyclase installs a novel carbon-carbon macrocycle into heptapeptides, expanding the diversity of plant-derived cyclic peptides. By interconverting two distinct cyclases through targeted mutations, we illuminate how these enzymes evolve new functions. This work highlights the evolutionary dynamics of plant peptide natural products and their potential applications in drug discovery and biotech crop development, while illustrating how genomic archaeology can reveal lost biosynthetic capabilities.

## Introduction

Plant peptides encompass diverse classes such as cyclotides (1), orbitides(2), defensins (3), thionins (4), snakins (5), and burpitides (6), each distinguished by unique structural features including disulfide bridges, cyclic backbones, and post-translational modifications. These peptides play important roles in plants’ adaptation to environmental niches by mediating pathogen and herbivore defense, growth regulation, stress responses, metal chelation, and root-microbiome interactions (6–8). Among these, burpitides are cyclic peptides characterized by distinct macrocycles installed by catalytic BURP domains specific to plants (6). Like other ribosomally synthesized and post-translationally modified peptides (RiPPs), their precursor peptides are genetically encoded and translated by the ribosome before undergoing class-defining post-translational modifications and subsequent proteolysis to produce mature peptides (9, 10). The BURP-domain proteins were initially named after their four founding members: BNM2, USP1, RD22, and PG1b, and contain a conserved sequence motif consisting of four sets of interspaced cysteine-histidine residues (11). Although not all BURPs demonstrate catalytic activity (12), this motif was recently shown to form a copper-dependent catalytic center that mediates various C-C, C-N and C-O bond formations necessary for burpitide macrocyclization (13, 14). While the biological functions of most burpitides remain elusive, they have been shown to exhibit analgesic (15, 16), tubulin-binding (14, 17, 18), and insecticidal properties (19, 20). The growing interest in peptide therapeutics highlights the importance of exploring new cyclic peptide scaffolds, such as those of burpitides, and macrocyclases key to cyclic peptide production that are otherwise chemically challenging or unfeasible (21, 22). The versatility of burpitide biosynthetic systems has also sparked interest in utilizing them as platforms for introducing engineered, genetically encoded agronomic traits (23).

A common feature of some BURP-containing proteins is a highly repetitive region at the N-terminus, which is often organized into strict tandem repeats with a specific pattern of core peptide motifs linked by recognition sequences (13, 24). Tandem repeats within genes are known to duplicate and reduce themselves through a process known as slipped strand mispairing (25–27). As tandem repeats are a common feature of many eukaryotic RiPPs, this dynamic may contribute to how core peptides and their repeat numbers are observed to vary (28). In the case of AhyBURP, an autocatalytic burpitide cyclase from the peanut plant (*Arachis hypogaea*), the enzyme’s turnover rate was shown to directly correlate with the number of core peptide motifs within the protein (13). The first burpitide precursor identified with a set of tandem repeats in this manner was LbaLycA, which produces lyciumin peptides with angiotensin-converting enzyme (ACE) inhibitory and renin inhibitory activities, found in goji berry (*Lycium barbarum*) (23). Lyciumins contain a macrocycle formed by linking the indole nitrogen of a tryptophan to the α-carbon of a glycine within the core peptide, and are produced by many members of the plant family *Solanaceae* (23). However, several lineages within *Solanaceae* appear to have lost the ancestral lyciumin precursor gene, an evolutionary pattern commonly observed in specialized metabolism (29, 30). Nicotiana benthamiana, for example, is commonly used for transient transgene expression (Fig. 1A) and is devoid of any lyciumin precursor gene homolog. It was therefore employed in a previous study as a convenient host for the heterologous reconstitution of LbaLycA (23).

**Figure 1.**
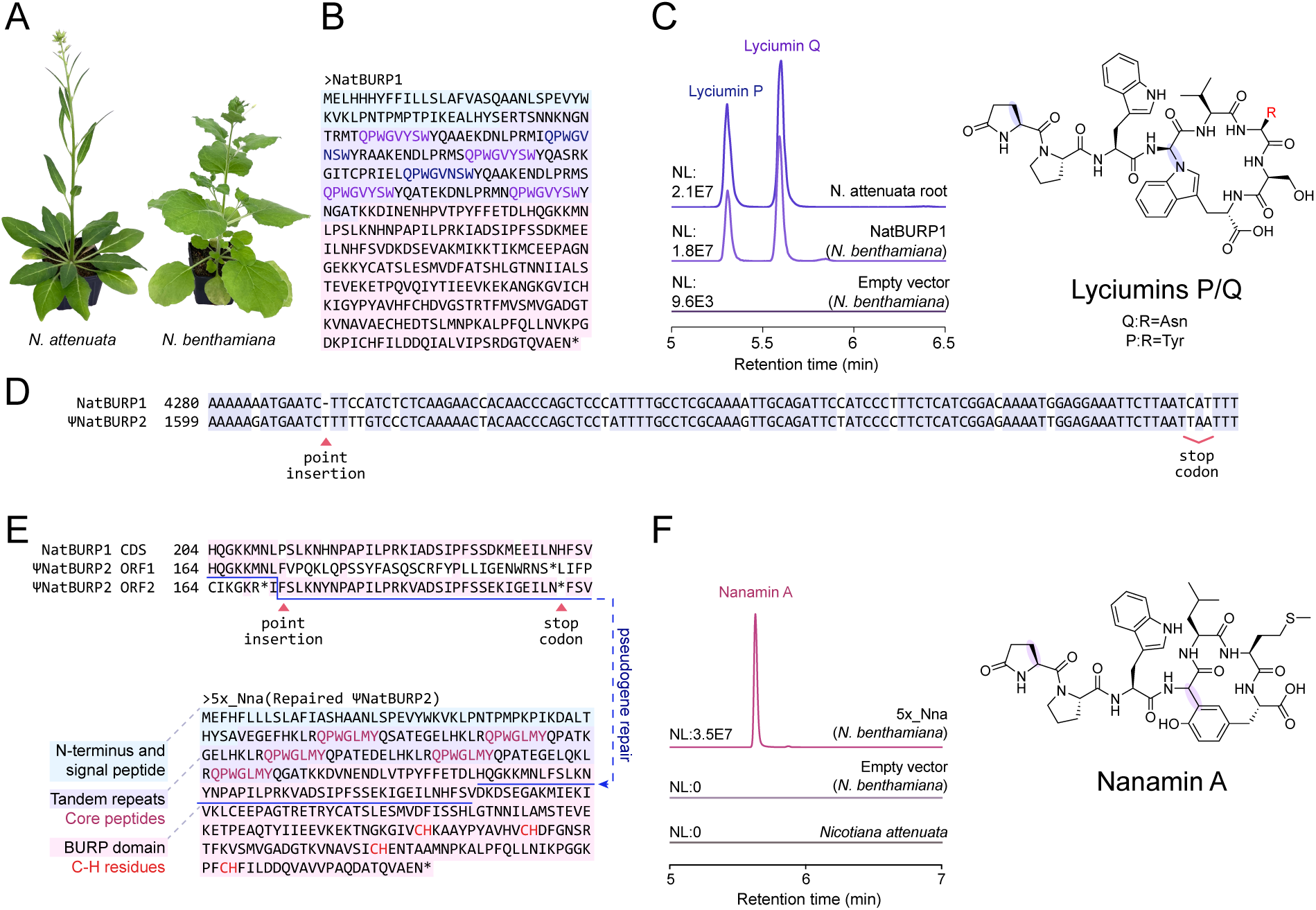
Characterization of lyciumin and nanamin biosynthesis in *Nicotiana*. (**A**) *N. benthamiana* (top) and *N. attenuata* (bottom). (**B**) Sequence of the lyciumin precursor gene from *N. attenuata*, *NatBURP1*. (**C**) Lyciumins P and Q from peptide extracts from *N. attenuata* vs. heterologous expression in *N. benthamiana*. Left: Combined EICs (954.41275, 1003.43084 m/z, 5 ppm) from LC-MS analysis of lyciumins P and Q. Axis indicates relative abundance. Right: structure of lyciumins P and Q, for which “R” indicates asparagine tyrosine residues, respectively. (**D**) Partial sequence alignment of *NatBURP1* and *ΨNatBURP2*. (**E**) Repair of the pseudogene *ΨNatBURP2*. By removing the point insertion and premature stop codons, we created the reconstituted gene *5x_Nna*. (**F**) Heterologous production of Nanamin A through transgenic expression of *5x_Nna*. Left: combined EICs (875.37564 m/z, 5 ppm) from LC-MS analysis. Right: structure of nanamin A. The macrocyclic bond and pyroglutamate are highlighted in purple.

Upon further investigation of the thirteen *Nicotiana* species with sequenced genomes, we found that only the coyote tobacco (*Nicotiana attenuata*) retains a direct homolog of the *LbaLycA*, namely *NatBURP1* (Fig. 1B). *N. attenuata* is a widely studied model organism in chemical ecology due to its complex interactions with herbivores, particularly in its ability to dynamically adjust its chemical defenses based on herbivore attack (31). In response to herbivory, *N. attenuata* can initiate several defensive maneuvers via the methyl jasmonate signaling system (32). These include altering its flowering timing to attract different pollinators (33), producing defense compounds or proteins including triterpenoids, caffeoylputrescine-green leaf volatile compound CPH, and trypsin proteinase inhibitors to deter or resist herbivory (34– 36), or attracting predatory insects to feed on its herbivores (37). These studies demonstrate how *N. attenuata* actively engages with and shapes its biotic environment in the struggle to survive and reproduce. While this chemical plasticity helps *N. attenuata* thrive in challenging ecosystems, maintaining such defenses also carries significant metabolic costs, and the evolutionary arms race ensures that not every adaptation will remain effective indefinitely (38– 40). The presence of a unique lyciumin precursor gene in *N. attenuata* raised intriguing questions about the evolutionary dynamics of cyclic peptides within *Nicotiana*, which prompted us to investigate its peptide products and their biosynthetic machinery.

## Results

### Cyclic peptides from *N. attenuata*

While lyciumins have been isolated from several *Solanaceae* species (23), there are no previous reports of their isolation from within the subfamily *Nicotianoideae*. We extracted tissue samples from *N. attenuata* and used liquid chromatography–mass spectrometry (LC-MS) to identify the presence of two distinct cyclopeptides within the roots, which we termed lyciumins P and Q (Fig. 1C). These chemotypes are consistent with those encoded by the gene *NatBURP1*, a BURP-domain protein from *N. attenuata*. The structure of this gene is similar to previously described genes encoding ‘fused’ burpitide cyclases, wherein a single polypeptide chain contains both the BURP domain and the core peptides (41). We confirmed the activity of this gene by cloning *NatBURP1* into the pEAQ-HT vector, transforming it into Agrobacterium tumefaciens LBA4404, then syringe-infiltrating the leaves of N. benthamiana (42–44), which resulted in the production of lyciumins P and Q (Fig. 1C).

NatBURP1 shares 90% and 83% amino acid sequence identity to LbaLycA and StuBURP, lyciumin precursors from *L. barbarum* and *Solanum tuberosum*, respectively (SI Appendix Fig. S1). This close homology suggests that the progenitor of these genes was most likely carried by the last common ancestor of these three species, which dates to approximately 24 million years ago (45). If lyciumin is indeed ancestral, it may explain why *N. benthamiana* can serve as a suitable host for its production, despite having lost its own lyciumin precursor gene.

Previous studies have shown that the BURP domain functions solely as a peptide cyclase, and that the full biosynthesis requires the activity of other enzymes (such as peptidases and glutaminyl transferases) free the core peptide through the proteolysis of the recognition sequence, and to convert N-terminal glutamine into pyroglutamate (23). While it was initially thought that nonspecific processive enzymes were responsible for completing the biosynthesis of lyciumin in *N. benthamiana*, it is also possible that, apart from the precursor protein itself, *N. benthamiana* has inherited specific biosynthetic machinery from an ancestor that once produced lyciumin.

While studying *NatBURP1*, we discovered another locus within the genome of *N. attenuata,* located just 0.8 Mbp away on the same chromosome. This newly identified locus encodes *ΨNatBURP2*, which shares 83% sequence identity to *NatBURP1* at the protein level, which suggests that the two are close homologs (SI Appendix, Fig. S1). However, *ΨNatBURP2* cannot function as a lyciumin precursor because it lacks the lyciumin motif [QPX₅W], where X represents any amino acid. Instead, *ΨNatBURP2* encodes several repeats of the core peptide [QPWGLMY], which is one amino acid shorter and contains a C-terminal tyrosine instead of tryptophan (Fig. 1). Nevertheless, *ΨNatBURP2* appears to be a pseudogenized locus due to a frameshift mutation and the presence of premature stop codons. (Fig. 1D).

### Resurrecting the pseudogene *ΨNatBURP2*

To investigate the original function of this pseudogene, we aligned its exons with homologous genes within Solanaceae to reconstruct a functional coding DNA sequence. The frameshift was corrected, and the two premature stop codons were replaced with homologous coding codons from *NatBURP1*. To address the challenges posed by repetitive regions during DNA synthesis, we reduced the number of tandem repeats from eleven to five, enabling the de novo synthesis of the resurrected gene called *5x_Nna* (Fig. 1E).

After cloning it into the pEAQ-HT vector the transient expression of *5x_Nna* in *N. benthamiana* produced an analyte matching the cyclic form of the peptide [QPWGLMY] (Fig. 1F). The mass of the compound indicates the loss of H_2_ and NH_3_ molecules, consistent with cyclization and pyroglutamate formation, which are known to occur during the biosynthesis of burpitides such as lyciumin and moroidin (14, 41). To elucidate the structure of this new peptide, termed “nanamin” (from ***N****icoti**ana***), we purified 8 mg of peptide from 200 g of *N. benthamiana* fresh leaf tissue infiltrated with the *5x_Nna* construct. During the purification process, the compound was observed to oxidize spontaneously, which we hypothesized to be due to the oxidation of methionine to methionine sulfoxide (46, 47). The oxidized peptide was purified and structurally elucidated, and was shown to contain three post-translational modifications, including oxidized methionine and N-terminal pyroglutamate as hypothesized (SI Appendix, Table S1 and Figs. S2-S6). The cyclic bond was revealed to be a link between the phenolic meta-carbon of the C-terminal tyrosine and the α-carbon of glycine (Fig. 1F), which classifies nanamin as a hibispeptin-type burpitide based on its C(sp^2^)-C(sp^3^) bond. While other members of this group are linked from sidechain to sidechain (6, 24, 48), nanamin’s cyclic bond instead involves the backbone α-carbon of its glycine residue. The activation of the α-carbon appears unique to the cyclic bonds of lyciumin (N-C) and nanamin (C-C), a similarity likely stemming from the common evolutionary origin of *NatBURP1* and *ΨNatBURP2*.

### Identification of species retaining the wild-type nanamin precursor gene

Consistent with the pseudogenization of *ΨNatBURP2*, tissue extracts of *N. attenuata* do not contain nanamin (Fig. 1F). To investigate whether any other *Nicotiana* species still retain the wild-type nanamin genotype and chemotype, we grew and analyzed four additional tobacco species: *N. clevelandii*, *N. quadrivalvis*, *N. pauciflora*, and *N. obtusifolia*. While *N. obtusifolia* is only distantly related to *N. attenuata*, the two species are believed to have hybridized approximately one million years ago, resulting in *N. clevelandii* as a hybrid descendant of both lineages (49, 50) (Fig. 2). Notably, we detected the presence of nanamin in root extracts of both *N. clevelandii* and *N. pauciflora*. In addition to nanamin A, we discovered that its oxidized form, which we have termed nanamin B, also occurs naturally within these two species (Fig. 2C).

**Figure 2.**
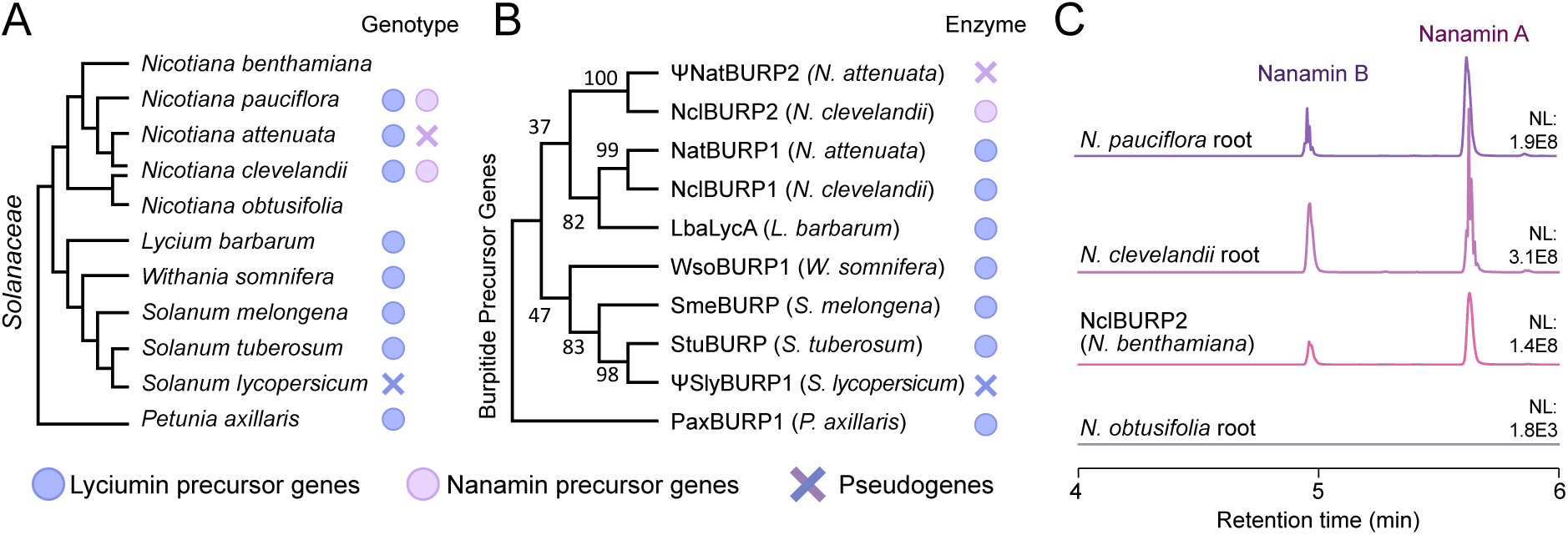
Phylogenetic and biochemical characterization of burpitide precursors in *Nicotiana*. (A) Phylogenetic tree of *Solanaceae* based on established taxonomic relationships. As a polyploid hybrid species, *N. clevelandii* is directly related to both *N. attenuata* and *N. obtusifolia*. (B) Maximum-likelihood phylogenetic analysis of burpitide precursor peptides. Bootstrap values based on 1000 replicates are shown at their respective nodes. (**C**) Combined EICs of nanamin A and B (875.37564, 891.37055 m/z, 5 ppm) from heterologous expression of *NclBURP2* in *N. benthamiana*, as well as root extracts of *Nicotiana* species.

Based on these leads, we used publicly available RNAseq datasets to assemble transcriptomes for *N. clevelandii* and *N. pauciflora*. From the assembled *N. clevelandii* transcriptome, we were able to identify both the full-length lyciumin and nanamin precursor genes, designated as *NclBURP1* and *NclBURP2*, respectively (SI Appendix, Table S2). We confirmed the activity of *NclBURP2* by transiently expressing it in *N. benthamiana* and found that it exhibited the same activity as *5x_Nna*. Although sequence reads supporting the presence of the nanamin precursor gene were detected in the transcriptome assembled for *Nicotiana pauciflora*, insufficient coverage prevented the reconstruction of the full-length sequence. This limitation was likely due to the absence of root tissue samples in the raw data. Therefore, our data show that the nanamin chemotype is retained by at least two close relatives of *N. attenuata*, where *NclBURP2* from *N. clevelandii* was characterized unequivocally as a nanamin precursor gene.

### The nanamin cyclase tolerates mutations within the core peptide

As the nanamin precursor gene encodes both the core peptide motif and a catalytic domain within the same polypeptide chain, we sought to evaluate the specificity of the nanamin cyclase itself by generating core peptide variants through mutagenesis. Using *5x_Nna* as the basis, we performed an alanine scan of the core peptide sequence [QPWGLMY]. By selectively mutating amino acids within this sequence to alanine, we were able to determine which residues were essential for cyclization and which were not. After infiltrating each construct into *N. benthamiana*, we did not observe the cyclization of the peptides [QPWALMY] and [QPWGLMA]. This indicates that both glycine and tyrosine are critical for product formation due to their participation in macrocyclization (Fig. 3A). The other 5 constructs produced cyclic peptides, including cyclo-[APWGLMY], which lacks N-terminal pyroglutamate (SI Appendix, Figs. S7-S15). This shows that the nanamin cyclase tolerates substitutions at non-critical sites within the core peptide.

**Figure 3.**
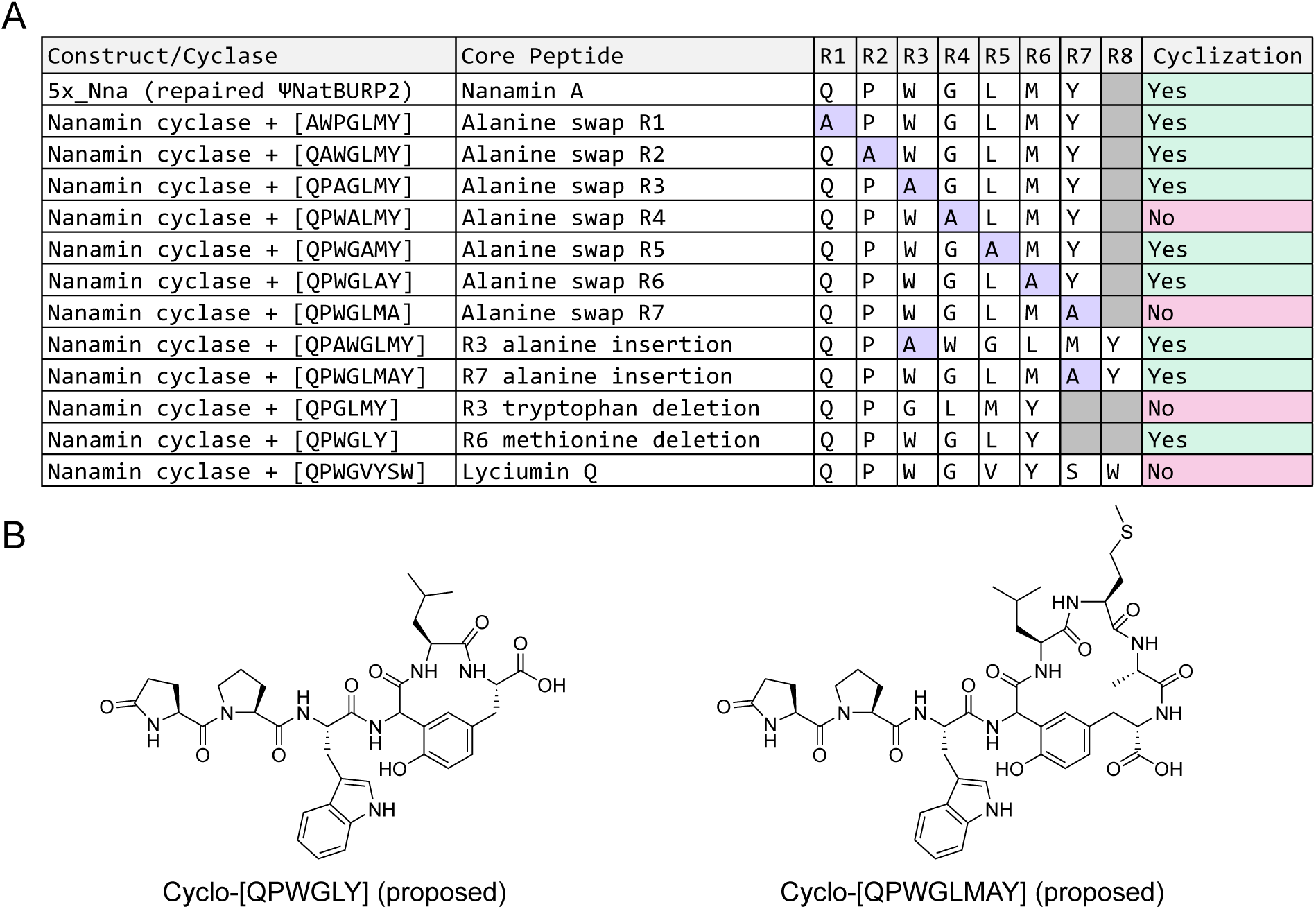
Substrate tolerance of the nanamin cyclase. (**A**) Cyclic peptide product formation by the nanamin cyclase in combination with various core peptide motifs. For each of the 13 constructs, different core peptides were paired with the repaired nanamin cyclase from *5x_Nna*. (**B**) Putative structures of cyclo-[QPWGLY] and [QPWGLMAY] based on MS/MS analysis.

Next, we tested constructs with either shortened or lengthened N- or C-terminal branches. We observed cyclic products for the peptides [QPWGLY], [QPAWGLMY], and [QPWGLMAY], but not [QPGLMY]. Analysis of MS/MS fragmentation patterns suggests cyclization occurs between tyrosine and glycine in all cases, even when the C-terminal branch was expanded and reduced (SI Appendix, Figs. S16-S18). The experiments demonstrate that the nanamin cyclase is remarkably tolerant of mutations within the core peptide.

### Molecular basis for the divergence of lyciumin and nanamin precursor genes

Maximum-likelihood phylogenetic analysis of select BURP-domain proteins from representative *Solanaceae* species suggests that nanamin precursor peptides evolved from an ancestral lyciumin precursor peptide, likely through gene duplication followed by neofunctionalization, which certainly occurred before the speciation of *N. attenuata* and *N. clevelandii* (Fig. 2B). To investigate the evolutionary path that led to the divergence of the lyciumin-producing *NatBURP1* and its nanamin-producing paralog, we examined the activities of several site-directed mutants of *NatBURP1* and *5x_Nna* using the heterologous expression system in *Nicotiana benthamiana* described previously. As with previous experiments, each mutant contained several copies of a specific core peptide attached to a cyclase.

When the nanamin cyclase from *5x_Nna* was paired with lyciumin core peptides, we did not detect the formation of lyciumin or any other cyclic peptide (Fig. 4B). However, when the lyciumin cyclase from *NatBURP1* was paired with nanamin core peptides, it produced two isomeric products: Nanamin A, and another cyclic peptide with the same mass but a distinct retention time (Fig. 4A). The fragmentation pattern of this isomer indicates that it contains a macrocyclic bond distinct from that of nanamin (SI Appendix, Fig. S19). While less efficient and specific, the lyciumin cyclase can accept the nanamin core peptide as its substrate. In contrast, the nanamin cyclase cannot accept lyciumin, but generates one major product from its native substrate (Fig. 4C). We hypothesize that specific adaptive mutations have occurred in an ancestral lyciumin cyclase, ultimately giving rise to the derived nanamin cyclase.

**Figure 4.**
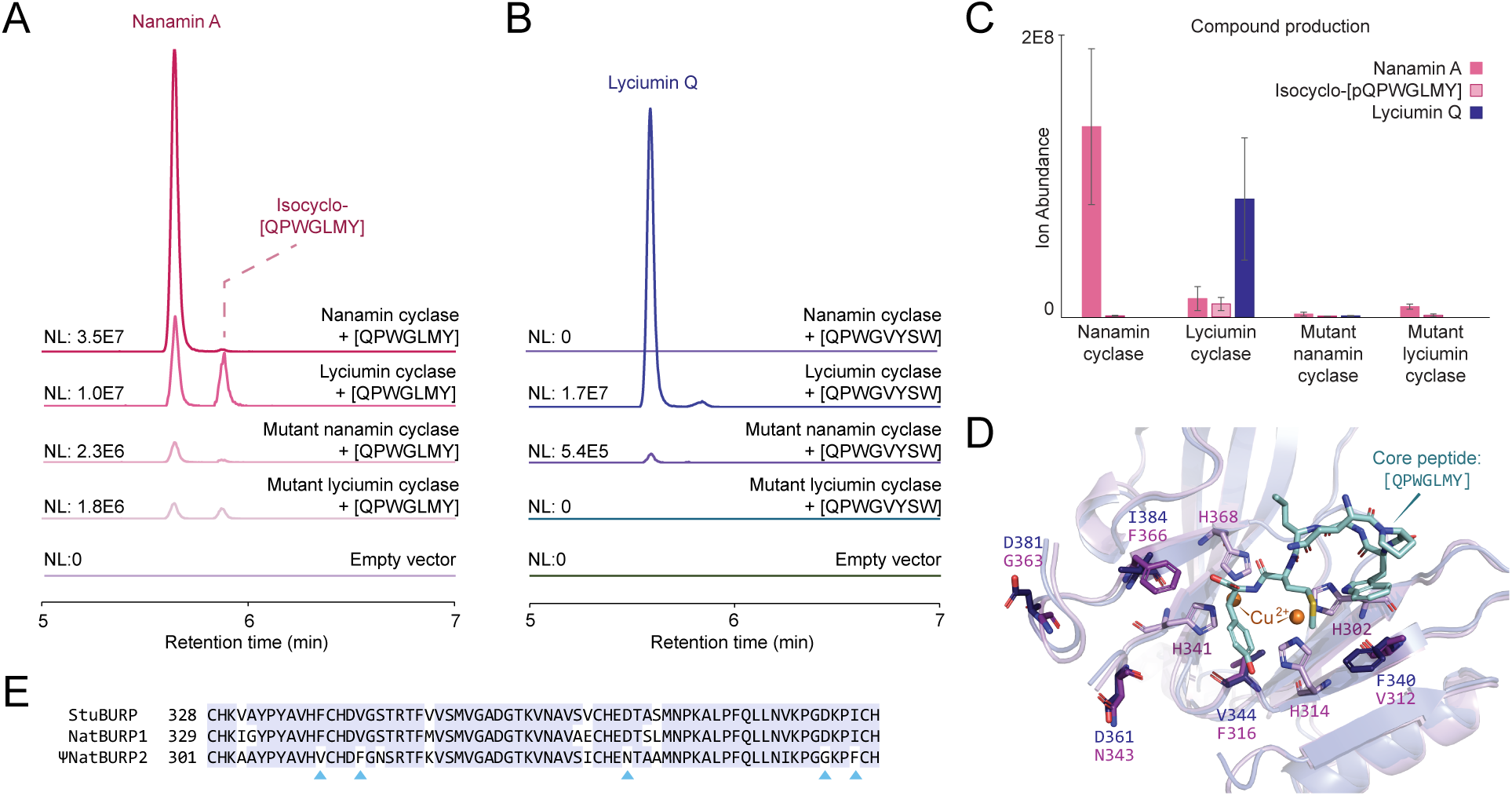
Cyclic peptide production from native and mutant BURP-domain macrocyclases *in vivo*. (**A**) Detection of cyclo-[QPWGLMY] in peptide extracts from infiltrated *N. benthamiana*. The constructs, containing nanamin core peptides paired with a catalytic BURP domain, were agroinfiltrated into *N. benthamiana*. MS/MS analysis indicates that the right-hand peak represents a nanamin isomer with a distinct macrocycle. (**B**) Detection of lyciumin Q in infiltrations of constructs containing lyciumin core peptides [QPWGVYSW] paired with catalytic BURP domains. (**C**) Cyclic peptide production organized by catalytic BURP domain. n = 3, error bars indicate ±1 sd. (**D**) Overlay of AlphaFold-predicted structures of NatBURP1 (blue) and 5x_Nna (purple), highlighting the active site. When included in the inputs, the model predicts the coordination of two copper ions by the central motif, along with the core peptide [QPWGLMY]. (**E**) Partial sequence alignment of NatBURP1, NatBURP2, and StuBURP, the lyciumin precursor from *Solanum tuberosum*. Mutant cyclases were generated by swapping the five residues marked by blue arrows between the catalytic domains of NatBURP1 and NatBURP2.

To identify the residues responsible for this functional transition, we generated structures for the two proteins using AlphaFold 3 (51) and aligned them (SI Appendix, Fig. S20). With accurate inputs, the structural models predicted the coordination of two copper ions in the center of the BURP domain and the localization of the core peptide [QPWGLMY] to the catalytic center (Fig. 4D). This structural comparison allowed us to identify five distinct residues in the vicinity of the active site (Fig. 4E). We then constructed two chimeric BURPs by swapping these residues between the two proteins (Fig. 4E). These mutant cyclases were paired with either the nanamin or lyciumin core peptides and expressed in *N. benthamiana* for functional testing. While both mutagenized cyclases retained the ability to produce nanamin *in vivo*, the mutagenized lyciumin cyclase lost its ability to produce lyciumin, while the mutagenized nanamin cyclase gained this ability (Fig. 4A, B). Although the catalytic efficiency of the mutants was reduced, we successfully interconverted the two peptide cyclases by swapping just five residues.

Given that the nanamin precursor gene is observed only in a few closely related *Nicotiana* species, the most parsimonious hypothesis is that it was derived from an ancestral lyciumin precursor within the *Nicotiana* genus (Fig. 2A, B). Examining the active site sequences of other homologs supports this hypothesis: at each of the five critical residues, *StuBURP* contains residues identical to *NatBURP1* and distinct from *ΨNatBURP2* (Fig. 4E). If *NatBURP1* more closely represents the ancestral state, its catalytic promiscuity suggests that *ΨNatBURP2* likely neofunctionalized substrate-first, through mutations that converted an ancestral lyciumin core peptide into a nanamin core peptide. Subsequent mutations in the BURP domain would have followed, harnessing the latent ancestral nanamin cyclization activity and enhancing this newly selected activity until the ancestral function was eventually lost. While the progenitor gene likely encoded multiple core peptides, the process of “repeat purification” could have homogenized these via cycles of duplication and reduction. Although *ΨNatBURP2* eventually became pseudogenized in *N. attenuata*, the gene and its associated chemotype have been retained in its close relatives, *N. clevelandii* and *N. pauciflora*.

## Discussion

The discovery of nanamins as unique, previously uncharacterized plant cyclic peptides, classified under the hibispeptin-type burpitides based on their C(sp^2^)-C(sp^3^) bond for macrocyclization, represent a significant addition to the chemical and biochemical diversity of plant peptides. Like other burpitides, nanamins are ribosomally synthesized and post-translationally modified to form macrocyclic bonds, but they display distinct structural features: a C-C bond between the phenolic meta-carbon of a conserved tyrosine and the α-carbon of a conserved glycine, and an unusually small ring size, distinguishing them from any previously described burpitides (6). This compact macrocycle, combined with N-terminal pyroglutamate formation and extensive sequence mutability at non-critical positions, makes nanamins particularly attractive for peptide engineering, alongside previously reported lyciumins (23), moroidins (14, 23), and cyclopeptide alkaloids (24). Collectively, these programmable peptides address key challenges in the development of peptide therapeutics, including proteolytic stability through cyclization and N-terminal protection, as well as potentially enhanced membrane permeability due to their compact structure and functional selection and refinement through million years of natural evolution. Furthermore, the nanamin cyclase’s ability to form challenging C-C bonds in small macrocycles—typically difficult to achieve through chemical synthesis— underscores the value of enzymatic approaches in accessing novel peptide scaffolds.

The serendipitous discovery of *ΨNatBURP2* joins a rich history of pseudogene studies that illuminate evolution in action. Classic examples include the L-gulono-γ-lactone oxidase (GLO) pseudogene in primates marking the loss of vitamin C synthesis after dietary changes (52), and rhodopsin pseudogenes in cave-dwelling fish, which document the loss of functional vision genes after moving to lightless environments (53). However, successfully resurrecting pseudogenes remains challenging, as deleterious mutations accumulate rapidly following the initial pseudogenization event. Mutations in the coding region, in particular, are expected to quickly narrow the paths for successful resurrection (54). Our relative ease in resurrecting *ΨNatBURP2* from *N. attenuata*, along with the preservation of functional nanamin biosynthesis in closely related *N. clevelandii* and *N. pauciflora*, provides a rare glimpse into an evolutionary transition in progress, showcasing the relatively recent gain and loss of a novel chemical trait in plants. In contrast to ancient protein resurrections, which typically reveal conservation of core mechanisms, our study demonstrates how novel enzymatic functions can emerge and disappear over relatively short evolutionary timescales in response to changing ecological environments.

Classical molecular evolution study of the glucocorticoid receptors showed that the evolutionary trajectory tends to be irreversible due to subsequent "restrictive" mutations that optimized its new specificity while making the ancestral conformation structurally incompatible, creating an epistatic ratchet that constrains evolutionary reversal (55). In contrast, studies of plant specialized metabolic enzymes such as sesquiterpene synthases reveal remarkable evolutionary plasticity, where even single mutations can alter product specificity and multiple mutation combinations can achieve similar functions (56). This flexibility likely reflects adaptation to fluctuating ecological pressures, where maintaining catalytic promiscuity and diverse product profiles provides selective advantages over strict functional optimization (57). In the case of nanamin evolution, the ancestral state represented by NatBURP1 is intrinsically promiscuous and can readily produce nanamin when provided with nanamin core peptide. The successful interconversion of lyciumin and nanamin cyclases by swapping just five active-site residues offers another example from plant specialized metabolic enzyme evolution that challenges the notion of irreversible protein evolutionary trajectories. The tolerance of the nanamin cyclase for a wide range of substrates, as well as the formation of two isomeric cyclopeptides by the lyciumin cyclase (Fig. 4A) provide compelling evidence that promiscuity and evolvability were likely selected traits in BURP-domain peptide macrocyclases. These features, in turn, offer unique biophysical and computational insights for the future design of natural-like biocatalysts with desirable functions and properties.

## Materials and Methods

### Plant Materials and Chemicals

*N. benthamiana* were originally gifted from the Lindquist laboratory at the Whitehead Institute. *N. attenuata* seeds were purchased from Grand Prismatic Seed. Seeds of *N. attenuata* (UT31X), *N. clevelandii* (TW30), *N. pauciflora*, and *N. obtusifolia* were received as a gift from Ian Baldwin and Wibke Seibt from the Max Planck Institute for Chemical Ecology, Jena, Germany. Plants were grown using a mixture of 3:1 potting soil to vermiculite (Lambert) with fertilizer, on a 16/8-hour day/night cycle. All chemicals and solvents used were purchased from Sigma-Aldrich.

### Construct design, synthesis, and cloning

Genes were either cloned from plant cDNA, or synthesized *de novo* from Twist Biosciences. Synthetic constructs were codon optimized to reduce repetitiveness and for optimal expression in *N. benthamiana*. These sequences were synthesized with 20 bp overhangs and cloned into the pEAQ-HT vector using Gibson assembly (NEBuilder hi-fi assembly kit, New England Biolabs) after AgeI/XhoI digestion and gel purification (42). Plasmids were transformed into *Escherichia coli* DH5a and purified for sequencing and transformation into *A. tumefaciens* LBA4404.

### Agrobacterium-mediated transient expression in *N. benthamiana*

Liquid cultures of *A. tumefaciens* LBA4404 were grown at 30 °C for 2 days at 250 rpm before collection by centrifugation. The cell pellets were resuspended in modified MM buffer [MgCl2 10 mM, MES 10mM, acetosyringone 100 µM, Tween 20 0.03% (v/v), Ascorbic acid 560 µM] at pH 5.6, normalized to an OD of 0.6 (44). *A. tumefaciens* suspensions were then syringe-infiltrated into the mature leaves of 4-to-5-week-old *N. benthamiana* plants, which were harvested and extracted after 7 days.

### Peptide extract preparation from plant tissue

To extract infiltrated *N. benthamiana* leaves and tissue samples from *N. attenuata* and *N. clevelandii*, 100 (±2) mg of tissue were measured and flash frozen with liquid nitrogen. Samples were lysed with 1 mL methanol and ∼10 zirconia beads (Research Products International) for 4 minutes at 30 hz with a Qiagen TissueLyser II. These samples were centrifuged before drying 800 µL of the clarified methanolic extract. The dried extracts were resuspended with 400 µL water and partitioned twice with n-hexane before further analysis.

### LC-MS analysis

All LC-MS analysis was conducted using a Vanquish Flex UHPLC system coupled with an Orbitrap Exploris 120 (Thermo Fisher scientific). Plant extracts were filtered with 0.2 µm PTFE syringeless filters (Whatman) before analysis. For reverse-phase LC separation analysis was performed with a Kinetex 2.6 µM, C18 100 Å, 150 x 3 mm^2^ column (Phenomenex). Solvent A was water with 0.1% formic acid, and solvent B was acetonitrile with 0.1% formic acid. Injections of 2µL were eluted at a flow rate of 0.5 mL/min with 5% B for 1 minute, 5-95% B for 7 minutes, 95% B for 3 minutes; 5% B for 4 minutes. MS analysis was performed in positive ion mode with a resolution of 60,000, RF lens of 70%, spray voltage of 3500 V, scan range of 300-1500 m/z, dd-MS2 (data-dependent MS/MS) taking the top 4 scans, dynamic exclusion of 0.5s, collision energy of 25 eV, and a scan width of 1. Data was collected and analyzed in Freestyle 1.8 (Thermo Fisher Scientific).

### Purification and structural elucidation of nanamins

To identify the structure of nanamin, 200 grams of *N. benthamiana* leaves infiltrated with the construct 5x_NnaBURP were used. The leaves were ground and extracted twice with 1 L methanol before drying and resuspending in 1L of water. The aqueous phase was washed twice with n-hexane before lyophilization, resuspension in methanol, and fractionation on a Sephadex LH-20 column (l=18 inches, i.d. = 1.5 inches) with methanol as the mobile phase. During the purification process, Nanamin A was observed to spontaneously oxidize into Nanamin B, forming methionine sulfoxide from methionine. This oxidation is well known to occur spontaneously (46, 47), and Nanamin B is also observed in extracts of *N. clevelandii* (Fig. 2C), we isolated the oxidized form to elucidate the structure of nanamin A. Fractions from the LH20 were concentrated and run on a Shimadzu semi-preparative HPLC (column: Kinetex® 5 μm C18 100 Å, 250x10 mm; mobile phase A: water + 0.1% formic acid, B: acetonitrile + 0.1 % formic acid, isocratic elution at 20% B, flow rate: 3 mL/min, injection: 1-15 mg dissolved in methanol and diluted in the starting conditions to 300 μL). Collection of the corresponding peak yielded 8 mg of Nanamin B. NMR spectra were recorded at 298 K on a 700 MHz Bruker AVANCE-Neo NMR spectrometer equipped with a cryoprobe. The compound was dissolved in DMSO-d_6_ (Cambridge Isotope Laboratories Inc.). Next to ^1^H-NMR spectra, DEPT-Q-^13^C, COSY, HSQC-DEPT, HMBC, and ROESY spectra were acquired. NMR data was analyzed using TopSpin 4.4.1. The residual solvent signals of DMSO (δ_H_ 2.50 ppm, δ_C_ 39.52 ppm) were used for referencing the 1D NMR spectra (58). The planar structure was identified through investigation of the acquired NMR spectra as the one of Nanamin B. Crucially, the signals were assigned to all predicted amino acids. This included the hypothesized O-methionine, the S-methyl group of which naturally overlapped in ^1^H-NMR (δ_H_ 2.50 ppm) with the structurally related solvent DMSO, however, was resolved in ^13^C NMR (δ_C_ 37.9 ppm). For, pyroglutamate while the ^13^C chemical shift of C-α (δ_C_ 53.7 ppm) and C-δ (δ_C_ 177.3 ppm) would only differ slightly between glutamate and pyroglutamate, HMBC correlations from C-α (δ_H_ 4.29 ppm) to C-δ (δ_C_ 177.3 ppm), as well as from pGlu-NH (δ_H_ 7.67 ppm) to all backbone carbons C-α (δ_C_ 53.7 ppm), C-β (δ_C_ 23.8 ppm), C-γ (δ_C_ 29.0 ppm), and C-δ (δ_C_ 177.3 ppm) clearly show the cyclic nature of the amino acid in the peptide. The third modification to the peptide regards the formation of the macrocycle. The presence of the Gly-C-α (δ_H_ 5.60 ppm, δ_C_ 50.6 ppm) as a CH group shows this as the point of cyclisation, whereas the relatively low ^13^C chemical shift excludes a C-O bond. Additionally, the absence of an AABB coupling pattern from tyrosine implies an asymmetrical substitution pattern. Finally, HMBC correlations from Gly-C-α (δ_H_ 5.60 ppm) to Tyr-C-ζ (δ_C_ 153.0 ppm), and somewhat weaker to the sp_2_-hybridized quaternary Tyr-C-ε (δ_C_ 122.8 ppm), which is connected directly to Gly-C-α. As the closure of the ring generates a new stereocenter, ROESY analysis was performed. However, most observed correlations were found within amino acids and no correlations to Gly-C-α were observed, indicating an orientation towards the Tyr-OH and not towards Tyr-H-δ. As the conformational change of tyrosine is independent of the configuration Gly-C-α, it was not assigned.

Next to the signals observed for the assigned structure, some signals are multiplied in ^1^H and ^13^C NMR (see related signals in table S1). In ^1^H NMR, signals in tryptophan from the aromatic NH (δ_H_ 10.81 and 10.73 ppm), H-ζ (δ_H_ 7.29 and 7.25 ppm), H-θ (δ_H_ 6.95 and 7.58 ppm), and the signal for H-α (δ_H_ 4.50 and 4.66 ppm) all show distinct chemical shifts and integrals implying a ratio of ca. 5:2 between the molecular species. In ^13^C NMR, next to the peak of the S-methyl group of O-methionine at 37.9 ppm, another peak appears at 38.2 ppm a smaller peak appears at 37.8 ppm. The same pattern appears with O-Met-C-γ (δ_C_ 49.5 ppm, smaller peak at 49.48 ppm and similar sized peak at 50.00 ppm). Also, each ^13^C signal of the Leu-CH_3_ groups appears as a “doublet” with a distance of 0.02 ppm between the peaks, and there are smaller twins of each doublet shifted from 21.8 ppm to 21.6 ppm for one methyl group, and from 22.7 to 22.8 ppm for the other. Additionally, for most signals from Trp, Pro, and Tyr, as well as for Gly, smaller twins can be observed in the spectrum. Due to the flexibility of the molecule and the limited signal recovered, it was not possible to find a definitive reason for this phenomenon. Likely explanations would be the presence of different conformers of Nanamin B, which has been described for other cyclic peptides (59). Additionally, non-specific oxidation of methionine generates two chemically distinct S- and R-diastereomers with potentially different chemical shifts (60), which likely accounts for chemical shift differences of carbons in proximity to the sulfur atom. Finally, if both configurations of glycine-tyrosine C-C bond would be generated by the cyclisation reaction, this could impact the chemical shifts of neighboring amino acids Trp, Tyr, Pro, and Leu. In conclusion, the structure of Nanamin B was established to be a burpitide of the hibispeptin-type, featuring a Tyr-C to C bond (6). Out of this class, nanamin is the first example that contains a Tyr-C to glycine C-α bond. While the principal structure of Nanamin B was confirmed, there are open questions regarding potential diastereomers from sulfur oxidation and from macrocycle formation at Gly-C-α that should be addressed in further work on this scaffold.

### Phylogenetic analysis

A phylogenetic analysis was performed using the protein sequences of ten lyciumin and nanamin precursor genes in *Solanaceae*. Tandem repeat regions were removed prior to sequence alignment with MUSCLE due to significant variation among the genes. The phylogenetic tree was constructed using MEGA 11 with the maximum-likelihood method, applying 1000 bootstrap replicates, the JTT model, gamma-distributed rates with invariant sites (5 discrete categories), and nearest neighbor interchange.

### *De novo* transcriptome assembly

Using sequence reads deposited at NCBI (Accessions SRR6918819 to SRR6918830 and SRR2913179 to SRR2913196), transcriptomes for *N. clevelandii* and *N. pauciflora* were assembled using the transXpress SnakeMake pipeline (61). While we successfully identified the lyciumin precursor gene in both species, we were only able to identify the full-length nanamin precursor gene in *N. clevelandii*. Sequence reads suggest that the nanamin precursor gene exists in *N. pauciflora* but is not sufficiently represented in the data to assemble the full-length gene. The sequences for *NclBURP1*, *NclBURP2*, and *NpaBURP1* will be deposited at NCBI upon publication of the paper.

### Molecular cloning

100 mg of root tissue was flash frozen and lysed in a Qiagen Tissuelyser II for 4 min at 30 hz with approximately 10 zirconia beads (2.3 mm, Research Products international). Total RNA was extracted following the protocol for the Qiagen RNeasy Plant Mini Kit. The extracted RNA was used to generate cDNA libraries with the Invitrogen Superscript III First-Strand Synthesis System using oligo(dT) primers. The genes were subsequently amplified using primers designed with overlapping ends complementary to the pEAQ-HT vector and amplified with Q5 polymerase. Following AgeI/XhoI restriction digestion, the genes were cloned into the vector backbone using the NEBuilder Hi-Fi Assembly Kit (New England Biolabs). *NatBURP1* was cloned using primers (NatBURP1-For, TATTCTGCCCAAATTCGCGAATGGAGTTGCATCACCATTACTTCT; NatBURP1-Rev, TGAAACCAGAGTTAAAGGCCCTAGTTCTCAGCCACTTGAGTTCCA) based on the *N. attenuata* UT31X genome (NIATv7_g20816/XP_019248232.1), while *NclBURP2* was cloned using primers (NclBURP2-For, TATTCTGCCCAAATTCGCGACTATTTAGTTCAGTTACAACATCAATG; NclBURP2-Rev, TGAAACCAGAGTTAAAGGCCTTGTGACTATTTCATAAAATGACTGAA), based on the assembled transcriptome. *NatBURP1* is available as A0A1J6ITY3 (Uniprot) and NIATv7_g20816 (*N. attenuata* data hub).

## Author Contributions

E.M.S. and J.-K.W. designed the research. E.M.S. performed most experiments. J.K.R. conducted peptide purification and NMR analyses. E.M.S. and J.-K.W. wrote the manuscript with inputs from J.K.R..

## Conflict of Interests

J.-K.W. is a member of the Scientific Advisory Board and a shareholder of DoubleRainbow Biosciences, Galixir, and Inari Agriculture, which develop biotechnologies related to natural products, drug discovery and agriculture. All other authors have no competing interests.

## Acknowledgements

This work was supported by the National Institute of Food and Agriculture of the United States Department of Agriculture, the Keck Foundation, the Schooner Foundation, and the Chan Zuckerberg Initiative Neurodegeneration Challenge Network. We would like to thank Dr. Ian Baldwin and Wibke Seibt of the Max Planck Institute for Chemical Ecology for generously providing *Nicotiana* seeds, as well as Matthew Hill and Erin Reynolds for their helpful advice and discussions. We would also like to extend special thanks to Stefano Rosa, Jennifer Sherk, Colin Kim, Corina Simian, Jason Guo, and the staff of the Northeastern University EXP building and the Whitehead Institute.

