## Supporting Information for "The emergence and loss of cyclic peptides in *Nicotiana* illuminate dynamics and mechanisms of plant metabolic evolution"

### Table of Contents


| | Title | Page |
|---|---|---|
| Table S1 | NMR data of nanamin B in DMSO-$d_6$ (700 MHz, 298 K) | 2 |
| Table S2 | Sequences of lyciumin and nanamin precursor genes from *Solanaceae* | 3 |
| Figure S1 | Alignments of lyciumin and nanamin precursor genes from *Solanaceae* | 5 |
| Figure S2 | $^1$H NMR spectrum of nanamin B (DMSO-$d_6$, 700 MHz, 298 K) | 6 |
| Figure S3 | $^{13}$C-DEPT-Q NMR spectrum of nanamin B (DMSO-$d_6$, 176 MHz, 298 K) | 7 |
| Figure S4 | COSY NMR spectrum of nanamin B (DMSO-$d_6$, 700 MHz, 298 K) | 8 |
| Figure S5 | HSQC-DEPT NMR spectrum of nanamin B (DMSO-$d_6$, 700 MHz, 298 K) | 9 |
| Figure S6 | HMBC NMR spectrum of nanamin B (DMSO-$d_6$, 700 MHz, 298 K) | 10 |
| Figure S7 | MS/MS analysis of lyciumin P | 11 |
| Figure S8 | MS/MS analysis of lyciumin Q | 12 |
| Figure S9 | MS/MS analysis of nanamin A | 13 |
| Figure S10 | MS/MS analysis of nanamin B | 14 |
| Figure S11 | MS/MS analysis of cyclo-[APWGLMY] | 15 |
| Figure S12 | MS/MS analysis of cyclo-[QAWGLMY] | 16 |
| Figure S13 | MS/MS analysis of cyclo-[QPAGLMY] | 17 |
| Figure S14 | MS/MS analysis of cyclo-[QPWGAMY] | 18 |
| Figure S15 | MS/MS analysis of cyclo-[QPWGLAY] | 19 |
| Figure S16 | MS/MS analysis of cyclo-[QPWGLY] | 20 |
| Figure S17 | MS/MS analysis of cyclo-[QPAWGLMY] | 21 |
| Figure S18 | MS/MS analysis of cyclo-[QPWGLMAY] | 22 |
| Figure S19 | MS/MS analysis of isocyclo-[QPWGLMY] | 23 |
| Figure S20 | AlphaFold-predicted structures for NatBURP1 and NatBURP2 | 24 |

### Table S1. NMR data of nanamin B in DMSO-$d_6$ (700 MHz, 298 K)

| | | δC | | δH in ppm, M | |
|---|---|---|---|---|---|
| | | major | related signals | major | related signals |
| pGlu | C=O | 170.5 | | | |
| | α | 53.67 | 53.71 | 4.29, m | |
| | β | 23.8 | | 2.15, m | |
| | | | | 1.78, m | |
| | γ | 29.0 | | 1.99, t (7.9) | |
| | δ | 177.3 | | | |
| | NH | | | 7.67, s | |
| Pro | C=O | 168.2 | | | |
| | α | 59.21 | 59.09 | 4.33, m | |
| | β | 28.8 | | 1.6, m | |
| | | | | 1.89, m | |
| | γ | 24.00 | 24.07 | 1.81, m | |
| | | | | 1.65, m | |
| | δ | 46.07 | 46.83 | 3.51, m | |
| | | | | 3.35, m | |
| Trp | C=O | 171.8 | | | |
| | α | 54.4 | | 4.50, m, I=0.75H | 4.66, m, I=0.30 H |
| | β | 27.4 | | 3.12, m | |
| | | | | 3.01, m | |
| | γ | 109.91 | 110.02 | | |
| | δ | 123.5 | | 7.14, m | |
| | NH | | | 10.81, br s, I=0.69H | 10.73, br s, I=0.29H |
| | ε | 136.0 | | | |
| | ζ | 111.20 | 111.24 | 7.29, br d (7.8), I=0.77H | 7.25, br d (8.1), I=0.30H |
| | η | 120.72 | 120.74 | 7.02, m | |
| | θ | 118.27 | 118.23 | 6.95, m | |
| | ι | 118.1 | | 7.48, d (7.8), I=0.74 H | 7.58, d, 7.6, I=0.31H |
| | κ | 127.31 | 127.24 | | |
| | NH | | | 6.77, m | |
| Gly | C=O | 170.5 | | | |
| | α | 50.6 | | 5.60, br d (6.4) | |
| | NH | | | 8.29, m | |
| Leu | C=O | 166.0 | | | |
| | α | 52.86 | 52.84 | 4.18, m | |
| | β | 40.41 | 40.43 | 1.50, m | |
| | | | | 1.60, m | |
| | γ | 24.25 | 22.30 | 1.59, m | |
| | δ | 22.65 | 22.67, 22.77, 22.79 | 0.86, m | |
| | ε | 21.80 | 21.82, 21.66, 21.61 | 0.83, m | |
| | NH | | | 8.29, m | |
| O-Met | C=O | 176.8 | | | |
| | α | 51.53 | 51.46, 51.59 | 4.30, m | |
| | β | 24.65 | 24.70 | 2.02, m | |
| | | | | 1.87, m | |
| | γ | 49.53 | 49.48 | 2.60, m | |
| | | | | 2.69, m | |
| | δ | 37.85 | 37.83, 38.20 | 2.50, m | |
| | NH | | | n.a. | |
| Tyr | C=O | 171.2 | | | |
| | α | 54.5 | | 4.10, m | |
| | β | 35.1 | | 2.92, m | |
| | | | | 3.04, m | |
| | γ | 128.21 | 128.33 | | |
| | δ | 129.4 | | 7.18, m | |
| | ε | 122.85 | 122.69 | | |
| | ζ | 153.06 | 153.10 | | |
| | η | 115.02 | 115.10 | 6.67, m | |
| | θ | 130.5 | | 6.92, m | |
| | NH | | | n.a. | |

Data are reported for the observed major molecular species. For $^{1}$H NMR signals, where a distinct corresponding peak were observed, signals and integrals (I) are reported for both molecular species. For $^{13}$C NMR signals, the observed major signals are reported with two along with the related signals.

### Table S2. Sequences of lyciumin and nanamin precursor genes from *Solanaceae*

| Gene | ID, Species, Database | Sequence |
|---|---|---|
| NatBURP1 | NIATv7_g20816/<br>XP_091248232.1<br>Nicotiana attenuata<br>NADH/NCBI | MELHHHYFFILLSLAFVASQAANLSPEVYWKVKLPNTPMPTPIKEALHYSERTS<br>NNKNGNTRMTQPWGVYSWYQAAEKDNLPRMIQPWGVNSWYQAAKENDLPRMSQP<br>WGVYSWYQAAEKDNLPRMIQPWGVNSWYRAAKENDLPRMSQPWGVYSWYQATEK<br>DNLPRMNQPWGVYSWYNGATKKDINENHPVTPYFFETDLHQGKKMNLPSLKNHN<br>PAPILPRKIADSIPFSSDKMEEILNHFSVDKDSEVAKMIKKTIKMCEEPAGNGE<br>KKYCATSLESMVDFATSHLGTNNIIALSTEVEKETPQVQIYTIEEVKEKANGKG<br>VICHKIGYPYAVHFCHDVGSTRTFMVSMVGADGTKVNAVAECHEDTSLMNPKAL<br>PFQLLNVKPGDKPICHFILDDQIALVIPSRDGTQVAEN |
| ΨNatBURP2 | XP_019259593.1<br>Nicotiana attenuata<br>NCBI | MIEKIVKLCEEPAGTRETRYCATSLESMVDFISSHLGTNNILAMSTEVEKETPE<br>AQTYIIEEVKEKTNGKGIVCHKAAYPYAVHVCHDFGNSRTFKVSMVGADGTKVN<br>AVSICHENTAAMNPKALPFQLLNIKPGGKPFCHFILDDQVAVVPAQDATQVAEN |
| 5x_Nna | Nicotiana attenuata | MEFHFLLLSLAFIASHAANLSPEVYWKVKLPNTPMPKPIKDALTHYSAVEGEFH<br>KLRQPWGLMYQSATEGELHKLRQPWGLMYQPATKGELHKLRQPWGLMYQPATED<br>ELHKLRQPWGLMYQPATEGELQKLRQPWGLMYQGATKKDVNENDLVTPYFFETD<br>LHQGKKMNLFSLKNYNPAPILPRKVADSIPFSSEKIGEILNHFSVDKDSEGAKM<br>IEKIVKLCEEPAGTRETRYCATSLESMVDFISSHLGTNNILAMSTEVEKETPEA<br>QTYIIEEVKEKTNGKGIVCHKAAYPYAVHVCHDFGNSRTFKVSMVGADGTKVNA<br>VSICHENTAAMNPKALPFQLLNIKPGGKPFCHFILDDQVAVVPAQDATQVAEN |
| NclBURP1 | Nicotiana<br>clevelandii<br>NCBI | MELHHHYFFILLCLAFVASQAANLSPEVYWKVKLPNTPMPTPIKEALHYSERTS<br>NNKNGNTRMTQPWGVFSWYRAAKENDLPRMSQPFGVFSWYRAAKENDLPRMSQP<br>FGVFSWYNGATKKDINENHPVTPYFFETDLHQGKKMNLPSLKNHNPAPILPRKI<br>ADSIPFSSDKMEEILNHFSVDRDSEVAKMIKKTIKMCEEPAGNGEKKYCATSLE<br>SMVDFATSHLGTNNIIALSTEVEKETPQVQIYTIEEVKEKANGKGVICHKIGYP<br>YAVHFCHDVGSTRTFMVSMVGADGTKVNAVAECHEDTSLMNPKALPFQLLNVKP<br>GDKPICHFILDDQIALVIPSRDGTHVAEN |
| NclBURP2 | Nicotiana<br>clevelandii<br>NCBI | MEFHFLLLSLAFIASHAANLSPEVYWKVKLPNTPMPKPIKDALTHYSAAEGELH<br>KLRQPWGLMYQPATEGELHKLRQPWGLIYQPATEGELHKLRQPWGLMYQPATED<br>ELHKLHQPWGLIYQPATKDELHKLRQPWGLMYQPATEDELHKLRQPWGLMYQPA<br>TEGELQKLRQPWGLMYQPATEDELHKLRQPWGLIYQPATEGELQKLRQPWGLMY<br>QPATEDELHKLRQPWGLIYQPATEGELQKLRQPWGLMYQPATEDELHKLRQPWG<br>LMYQGATKKDVNENDLVTPYFFETDLHQGKKMNLLSLKNYNPAPILPRKVADSI<br>PFSSEKIGEILSQFSIDKDSEGAKMIEKIVKLCEEPAGTRETRYCATSLESMAD<br>FISSHLGTNNILAMSTEVEKETPEAQTYIIEEVKEKTNGKGIVCHKAAYPYAVH<br>VCHDFGNSRTFKVSMVGADGTKVNAVSICHENTAAMNPKALPFQLLNIKPGGKP<br>FCHFILDDQVALVPAQDATQVAEN |
| NpaBURP1 | Nicotiana<br>pauciflora<br>NCBI | MELHHHYFFILLFLAFVASQAANLSPEVYWKVKLPNTPMPTPIKEALHYSERTS<br>NNKNGNTIMPQPWGVNSWYQAAEKDNLPRMSQPWGVYSWYQAAKEKDLPRMSQP<br>WGVYSWYRAAKEKDLPRMSQPWGVYSWYQAAEKDNLPRMSQPWGVYSWYNGATK<br>KDINENLPVTPYFFETDLHQGKKMNLPSLKNHNPAPILPRKIADSIPFSSDKME<br>EILNHFSVDKDSEVAKMIKKTIKMCEEPAGNGEKKYCATSLESMVDFATSHLGT<br>NNIIALSTEVEKETPQVQIYTIEEVKEKANGKGVICHKIGYPYAVHFCHDVGST<br>RTFMVSMVGADGTKVNAVAECHEDTSLMNPKALPFQLLNVKPGDKPICHFILDD<br>QISLVIPSRDGTQVAEN |
| LbaLycA | AYN06992.1<br>Lycium barbarum<br>NCBI | MELHHHYFFILLSLAFIASHAANLSPEVYWKVKLPNTPMPRPIKDALHYSEASE<br>GDVHKLRQPWGVGSWYQAANEGDIKKLRQPYGVGIWYQAANEGDVKKLRQPWGV<br>GSWYQAANEGDVKKLRQPWGVGSWYQAANEGDVKKLRQPWGVGSWYQAANEGDA<br>NEGDVKKLRQPYGVGIWYQAANEGDVKKLRQPWGVGSWYQAANEGDVKKLRQPW<br>GVGSWYQAANEGDVKKLHQPWGVGSWYQAANEGDVKKLPQPWGVGSWYQAANEG<br>DVKKLRQPYGVGIWYEAANEGQVKKLRQPYGVGSWYNTATKKDVNENLPVTPYF<br>FETDLHQGKKMNLPSLKNYNPAPILPRKVADSIPFSSDKIEEILKHFSIDKDSE<br>GAKMIKKTIKMCEEQAGNGEKKYCATSLESMVDFTSSYLGTNNIIALSTLVEKE<br>TPEVQIYTIEEVKEKANGKGVICHKVAYPYAIHYCHSVGSTRTFMVSMVGSDGT<br>KVNAVSECHEDTAPMNPKALPFQLLNVKPGDKPICHFILDDQIALVPSQDATQV<br>SEN |

**Table S2 (continued)**

| Gene | ID, Species, Database | Sequence |
|---|---|---|
| WsoBURP1 | Withania somnifera<br>NCBI | MELHHHYFFILLSVTFVASDAANLSPEVYWKVKLANTPMPRPIKDALRNPAAQSDLHKLRQPWGVGSSYKPTTEDEVSKLRQPYGVYGWYQPAAEGDLHKLRQPWGVGSWYKPTTEVRFANYVSRLECMDGINQQPRVTFTNYANHGEWVPGINQPRRVRFANYVSRLECMDGINQQPRVTFTNYANHGEWVPGINQPRRVSFTNYVNHLECMYGINQPLRVTFTHYANHGEWVRGINQPLRDSEGANMIMKTIKMCEDPAGNGEEKYCATSLESMVDFTSSHLGTNKISAMSTEVEKETPEVQTYTIEEVKEKANGKGLICHKVAYPYAVHFCHDVGSTRTFMVSMVGADGTKVNAVSVCHEDTASMNPKALPFQLLNVKPGDKPICHFILDDQIALVPSQEATQVAKN |
| SmeBURP | Smechr0801593.1<br>Solanum melongena<br>Sol genomics network | MELHHQYYFFTLFSLVLVASQAANLSPEVYWKVKLPNTPMPKPIKDALHISEKTAYNGDKSTKISQPWGVGSWYQAAPENELHKVRQPWGVLGWYHDAPENELHKLRQPWGVGSWYQAALENELHRVRQPWGVLGWYHDAPENELHKLRQPWGVGSWYQAAPENELHRVRQPWGVLGWYHDAPENDLHKLRQPWGVGSWYNGVAKKDQHENHLVTPYFFETDLHQGKTMNLLSLKNYNPAPILPRKVVDSIPFSSEKIEEILSHFSADKDSERAEMIKKTIKMCEDPAGNGEVKHCATSLESMLDFTVSHLGTNNIIAISTEVEKETPEVQTYTIEKVEEKANGKGVVCHKVAYPYSVHFCHDVGSTRTFMVSMVGADGTKVNAVSVCHEDTAPMNPKALPFQLLNVKPGDKPICHFTLDDQIALFPSPNVPLQVTKN |
| StuBURP | AYN06994.1<br>Solanum tuberosum<br>NCBI | MELLHQYYFFTFFSVIFVVSHAANLSPEVYWRVKLPNTPMPTPIKDALHISEKTAYNGDGNTKISQPYGVFAWYQAASENELHKVRQPYGVDGWYKAASENELHKVRQPYGVFAWYKAITENELHKVRQPYGVFAWYKAATENELHKVRQPYGVFAWYKAASENVLHKVRQPYGVFAWYNDAAKKDLNDNHPVTPYFFETDLHQGKKMNLQSLKNYNPAPILPRKVVDSIAFSSDKIEEILNHFSVDKDSERAKDIKKTIKMCEEPAGNGEVKHCATSLESMIDFTLSHLGTNNIVAISTEVDKETPEVQTYTIEKVEEKANGKGVVCHKVAYPYAVHFCHDVGSTRTFVVSMVGADGTKVNAVSVCHEDTASMNPKALPFQLLNVKPGDKPICHFTLDDQIALFPSQNALLQVAEN |
| ΨSlyBURP1 | Solyc08g068140.4.1<br>Solanum lycopersicum<br>Sol genomics network | MELYHNQYYVFTIFSVIFVVSHAANLSPEVYWRVKFPNTPMPTPIKDALHINEKPAYNGDGNTKISQPWGVGAWYKAAPEDELHKIRQPWGVGAWYKAAPEDELHKIRQPWGVGAWYKAAPEDELHKIRQPWGVGAWYKVAPEDELHKIRQPWGVGAWYKAAPEDELHKIRQPWGVGAWYKAAPEDELHKIRQPWGVGAWYKAAPEDELHKIRQPWGVGAWYKAVPEDELHKIRQPWGVGAWYKAVPEDELHKIRQPWGVGAWYKAAPEDELHKIRQPWGVGAWYKAAPEDELHKIRQPWGVYRWYQAAPEDELHKIRQPYGVYRWYQAAPEDELHKIRQPYGVYRWYQAAPEDELHKIRQPYGVYSWYQAAPEDELHKIRQPYGVYSWYQAAPENELHKVHQPWGVGSWYNDAAKKDLNDNHPVTPYFFETDLHQGKKMNLEVSQKLQSSTHFATQSCRFNRFLIRQN |
| | Solyc08g068130.1.1 | TLKSLKNYNPAPILPRKVVDSIAFSSDKIEEILNHFSADKDSERAKDIKKTIKMCEEPAGNGEVKHCATSLESMIDFTLSHLGTNKIIAISTEVEKETPEVQTYTIEKVEEKANGKGVVCHKVAYPYAVHFCHDVGSTRTFMVSMVGADGTKVNAVSVCHEDTASMNPKALPFQLLNVKPGDKPICHFTLDDQIALFPSQNTDLQVAEN |
| PaxBURP1 | Peaxi162Scf00799g00412.1<br>Petunia axillaris<br>Sol genomics network | MKSQLLYSLTIFWLAFVASHAAISPEIYWKVKLPNTPIPKVIKDFLPLAVNEVPELKQDKLASKEKVFSGLQQPYGVFAWHRAATEDELHELQKDNAASKKKVFSGLQQPYGVFAWHRAATEDELHELQKHSTGSMEKVYYGLHQPYGVFAWHRAATEDELHEMQKDNTGSMEKVYYGLHQPFGVFAWHRAATGDEVRELKKQSPSINHVTAMNELQQLTERSNAKNDLNDKFLYKPYFLEEALEKGKIINFPSLKNKNEAPFLSRQFVESIPFSSQKIPEILKYLSIDSSSSKDAQKIEETIKLCEKPEIKHKEKKRCATSLESMVDFSISVLGTNNIKALTTEVAGETQMMQKYTIEEVQQVAEGDNMVCHKLSYAYAVHYCHVGGATKTYMVSMVGADGTKVKAVSVCHKDTSFWNPKGLPFVVLKVKPGTTPICHFLQDDQIVFVPAKEATKISDN |

A

```
 PaxBURP1     1 MKSQL--LYSLTIFWLAFVASHAA-ISPEIYWKVKLPNTPIPKVIKDFL---PLKFLYKPYFLEEALEKGKIINFPSLKNKNEAPFLSRQFVESI
ΨNatBURP2     1 MEFH------FLLLSLAFIASHAANLSPEVYWKVKLPNTPMPKPIKDALTHYSANLPVTPYFFETDLHQGKKMNLFSLKNYNPAPILPRKVADSI
 NclBURP2     1 MEFH------FLLLSLAFIASHAANLSPEVYWKVKLPNTPMPKPIKDALTHYSANDLVTPYFFETDLHQGKKMNLLSLKNYNPGPILPRTVADSI
 SmeBURP      1 MELHH-QYYFFTLFSLVLVASQAANLSPEVYWKVKLPNTPMPKPIKDAL-HIS-NHLVTPYFFETDLHQGKTMNLLSLKNYNPAPILPRKVVDSI
ΨSlyBURP1     1 MELYHNQYYVFTIFSVIFVVSHAANLSPEVYWRVKFPNTPMPTPIKDAL-HIN-NHPVTPYFFETDLHQGKKMNLESLKNYNPAPILPRKVVDSI
 StuBURP      1 MELLH-QYYFFTFFSVIFVVSHAANLSPEVYWRVKLPNTPMPTPIKDAL-HIS-NHPVTPYFFETDLHQGKKMNLQSLKNYNPAPILPRKVVDSI
 WsoBURP1     1 MELHH--HYFFILLSVTFVASDAANLSPEVYWKVKLANTPMPRPIKDAL-R-N-NHLITPYFFETDLHQGKKLNLPSLKNYNPAPFLPRKVAHSI
 LbaLycA      1 MELHH--HYFFILLSLAFIASHAANLSPEVYWKVKLPNTPMPRPIKDAL-HYS-NLPVTPYFFETDLHQGKKMNLPSLKNYNPAPILPRKVADSI
 NclBURP1     1 MELHH--HYFFILLCLAFVASQAANLSPEVYWKVKLPNTPMPTPIKEAL-HYS-NDLVTPYFFETDLHQGKKMNLLSLKNYNPAPILPRKIADSI
 NatBURP1     1 MELHH--HYFFILLSLAFVASQAANLSPEVYWKVKLPNTPMPTPIKEAL-HYS-NLPVTPYFFETDLHQGKKMNLPSLKNHNPAPILPRKIADSI
 NpaBURP1     1 MELHH--HYFFILLFLAFVASQAANLSPEVYWKVKLPNTPMPTPIKEAL-HYS-NLPVTPYFFETDLHQGKKMNLPSLKNHNPAPILPRKIADSI

 PaxBURP1    90 PFSSQKIPEILKYLSIDSSSSKDAQKIEETIKLCEKPEIKHKEKKRCATSLESMVDFSISVLGTNNIKALTTEVAGETQMMQKYTIEEVQQVAEG
ΨNatBURP2    90 PFSSEKIGEILNHFSVDKDSEG-AKMIEKIVKLCEEPA-GTRETRYCATSLESMVDFISSHLGTNNILAMSTEVEKETPEAQTYIIEEVKEKTNG
 NclBURP2    90 PFSSEKIGEILSQFSIDKDSEG-AKMIEKIVKLCEEPA-GNGEVRYCATSLESMADFISSHLGTNNILAMSTEVEKETPEAQTYIIEEVKEKTNG
 SmeBURP     93 PFSSEKIEEILSHFSADKDSER-AEMIKKTIKMCEDPA-GNGEVKHCATSLESMLDFTVSHLGTNNIIAISTEVEKETPEVQTYTIEKVEEKANG
ΨSlyBURP1    94 AFSSDKIEEILNHFSADKDSER-AKDIKKTIKMCEEPA-GNGEVKHCATSLESMIDFTLSHLGTNKIIAISTEVEKETPEVQTYTIEKVEEKANG
 StuBURP     93 AFSSDKIEEILNHFSVDKDSER-AKDIKKTIKMCEEPA-GNGEVKHCATSLESMIDFTLSHLGTNNIVAISTEVDKETPEVQTYTIEKVEEKANG
 WsoBURP1    91 PFSSDKIDEILNQFSIDKDSEG-ANMIMKTIKMCEDPA-GNGEEKYCATSLESMVDFTSSHLGTNKISAMSTEVEKETPEVQTYTIEEVKEKANG
 LbaLycA     92 PFSSDKIEEILKHFSIDKDSEG-AKMIKKTIKMCEEQA-GNGEKKYCATSLESMVDFTSSYLGTNNIIALSTLVEKETPEVQIYTIEEVKEKANG
 NclBURP1    92 PFSSDKMEEILNHFSVDRDSEV-AKMIKKTIKMCEEPA-GNGEKKYCATSLESMVDFATSHLGTNNIIALSTEVEKETPQVQIYTIEEVKEKANG
 NatBURP1    92 PFSSDKMEEILNHFSVDKDSEV-AKMIKKTIKMCEEPA-GNGEKKYCATSLESMVDFATSHLGTNNIIALSTEVEKETPQVQIYTIEEVKEKANG
 NpaBURP1    92 PFSSDKMEEILNHFSVDKDSEV-AKMIKKTIKMCEEPA-GNGEKKYCATSLESMVDFATSHLGTNNIIALSTEVEKETPQVQIYTIEEVKEKANG

 PaxBURP1   185 DNMVCHKLSYAYAVHYCHVGGATKTYMVSMVGADGTKVKAVSVCHKDTSFWNPKGLPFVVLKVKPGTTPICHFLQDDQIVFV-PAKEAT-KISDN
ΨNatBURP2   183 KGIVCHKAAYPYAVHVCHDFGNSRTFKVSMVGADGTKVNAVSICHENTAAMNPKALPFQLLNIKPGGKPFCHFILDDQVAVV-PAQDAT-QVAEN
 NclBURP2   183 KGIVCHKAAYPYAVHVCHDFGNSRTFKVSMVGADGTKVNAVSICHENTAAMNPKALPFQLLNIKPGGKPFCHFILDDQVALV-PAQDAT-QVAEN
 SmeBURP    186 KGVVCHKVAYPYSVHFCHDVGSTRTFMVSMVGADGTKVNAVSVCHEDTAPMNPKALPFQLLNVKPGDKPICHFTLDDQIALF-PSPNVPLQVTKN
ΨSlyBURP1   187 KGVVCHKVAYPYAVHFCHDVGSTRTFMVSMVGADGTKVNAVSVCHEDTASMNPKALPFQLLNVKPGDKPICHFTLDDQIALF-PSQNTDLQVAEN
 StuBURP    186 KGVVCHKVAYPYAVHFCHDVGSTRTFVVSMVGADGTKVNAVSVCHEDTASMNPKALPFQLLNVKPGDKPICHFTLDDQIALF-PSQNALLQVAEN
 WsoBURP1   184 KGLICHKVAYPYAVHFCHDVGSTRTFMVSMVGADGTKVNAVSVCHEDTASMNPKALPFQLLNVKPGDKPICHFILDDQIALV-PSQEAT-QVAKN
 LbaLycA    185 KGVICHKVAYPYAIHYCHSVGSTRTFMVSMVGSDGTKVNAVSECHEDTAPMNPKALPFQLLNVKPGDKPICHFILDDQIALV-PSQDAT-QVSEN
 NclBURP1   185 KGVICHKIGYPYAVHFCHDVGSTRTFMVSMVGADGTKVNAVAECHEDTSLMNPKALPFQLLNVKPGDKPICHFILDDQIALVIPSRDGT-HVAEN
 NatBURP1   185 KGVICHKIGYPYAVHFCHDVGSTRTFMVSMVGADGTKVNAVAECHEDTSLMNPKALPFQLLNVKPGDKPICHFILDDQIALVIPSRDGT-QVAEN
 NpaBURP1   185 KGVICHKIGYPYAVHFCHDVGSTRTFMVSMVGADGTKVNAVAECHEDTSLMNPKALPFQLLNVKPGDKPICHFILDDQISLVIPSRDGT-QVAEN
```

B

| | | | | | | | | | | | |
|---|---|---|---|---|---|---|---|---|---|---|---|
| 1: | PaxBURP1 | 100.00 | 56.09 | 56.09 | 57.66 | 56.20 | 56.57 | 58.76 | 59.12 | 58.03 | 58.03 | 58.03 |
| 2: | ΨNatBURP2 | 56.09 | 100.00 | 95.27 | 76.92 | 76.56 | 78.02 | 79.41 | 82.78 | 79.49 | 80.95 | 80.22 |
| 3: | NclBURP2 | 56.09 | 95.27 | 100.00 | 78.39 | 76.19 | 77.29 | 80.15 | 82.05 | 79.49 | 79.12 | 78.39 |
| 4: | SmeBURP | 57.66 | 76.92 | 78.39 | 100.00 | 88.89 | 88.89 | 83.33 | 83.39 | 82.67 | 82.67 | 81.95 |
| 5: | ΨSlyBURP1 | 56.20 | 76.56 | 76.19 | 88.89 | 100.00 | 95.34 | 83.33 | 82.67 | 81.23 | 82.31 | 81.59 |
| 6: | StuBURP | 56.57 | 78.02 | 77.29 | 88.89 | 95.34 | 100.00 | 82.97 | 83.39 | 81.95 | 83.03 | 82.31 |
| 7: | WsoBURP1 | 58.76 | 79.41 | 80.15 | 83.33 | 83.33 | 82.97 | 100.00 | 86.59 | 83.70 | 84.42 | 83.70 |
| 8: | LbaLycA | 59.12 | 82.78 | 82.05 | 83.39 | 82.67 | 83.39 | 86.59 | 100.00 | 88.09 | 89.89 | 89.17 |
| 9: | NclBURP1 | 58.03 | 79.49 | 79.49 | 82.67 | 81.23 | 81.95 | 83.70 | 88.09 | 100.00 | 97.48 | 97.12 |
| 10: | NatBURP1 | 58.03 | 80.95 | 79.12 | 82.67 | 82.31 | 83.03 | 84.42 | 89.89 | 97.48 | 100.00 | 99.28 |
| 11: | NpaBURP1 | 58.03 | 80.22 | 78.39 | 81.95 | 81.59 | 82.31 | 83.70 | 89.17 | 97.12 | 99.28 | 100.00 |

**Figure S1. Sequence alignments of lyciumin and nanamin precursor proteins from *Solanaceae*.** (**A**) Sequences aligned using MUSCLE. The tandem repeat regions were removed from each sequence before alignment to improve data analysis. (**B**) Percent identity matrix.

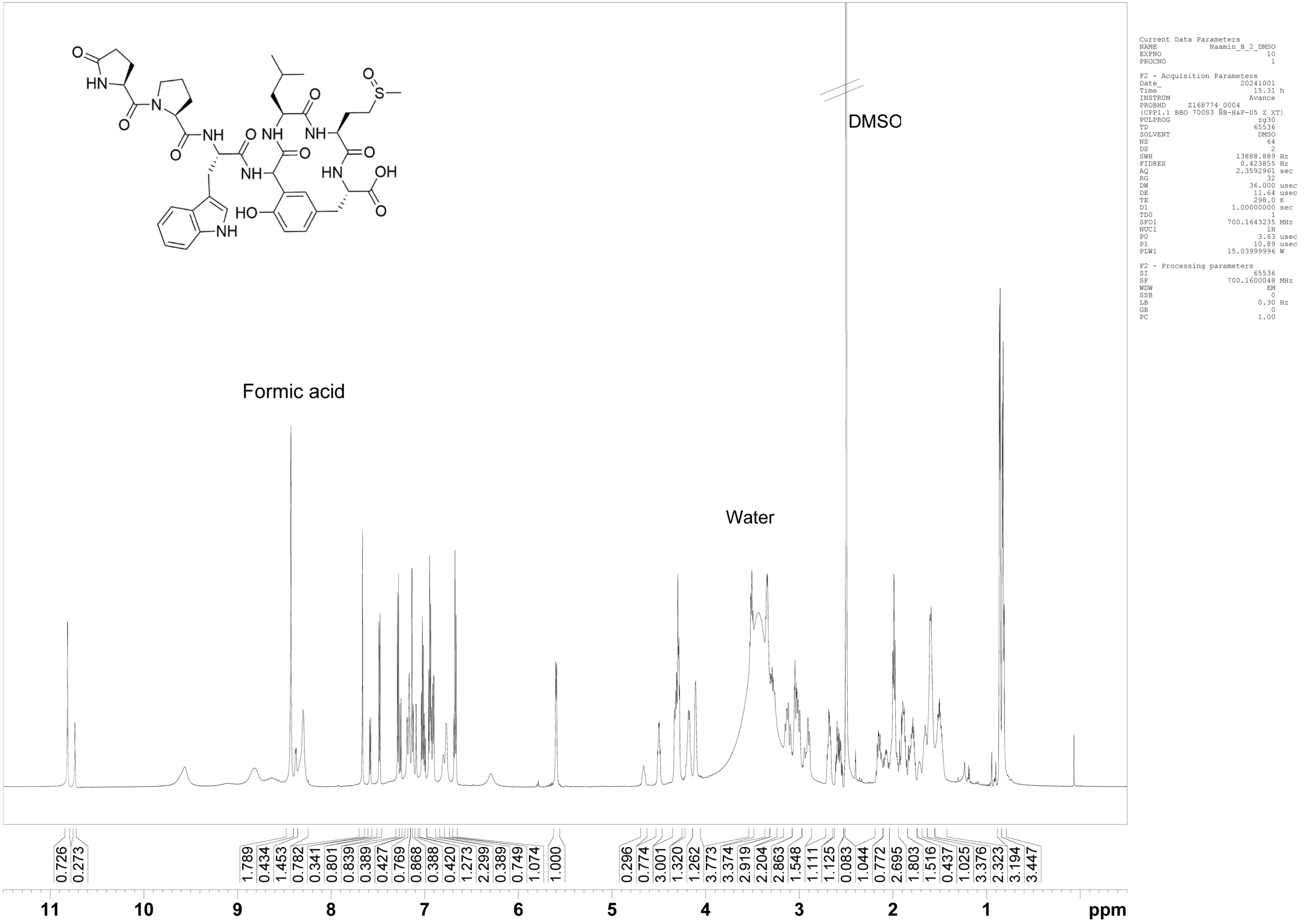


**Figure S2. $^1$H NMR spectrum of nanamin B (DMSO-$d_6$, 700 MHz, 298 K)**

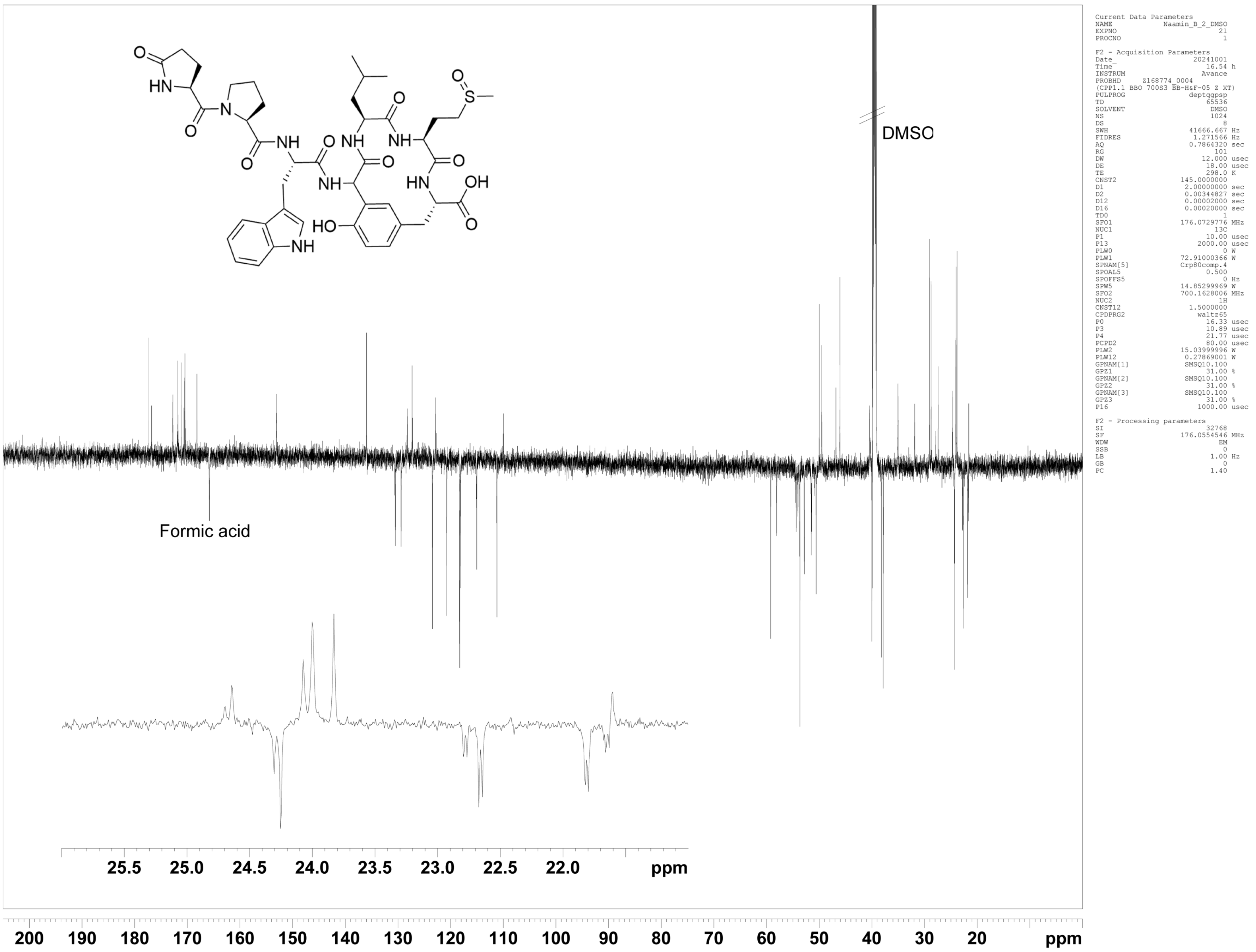


**Figure S3. $^{13}$C-DEPT-Q NMR spectrum of nanamin B (DMSO-$d_6$, 176 MHz, 298 K)**

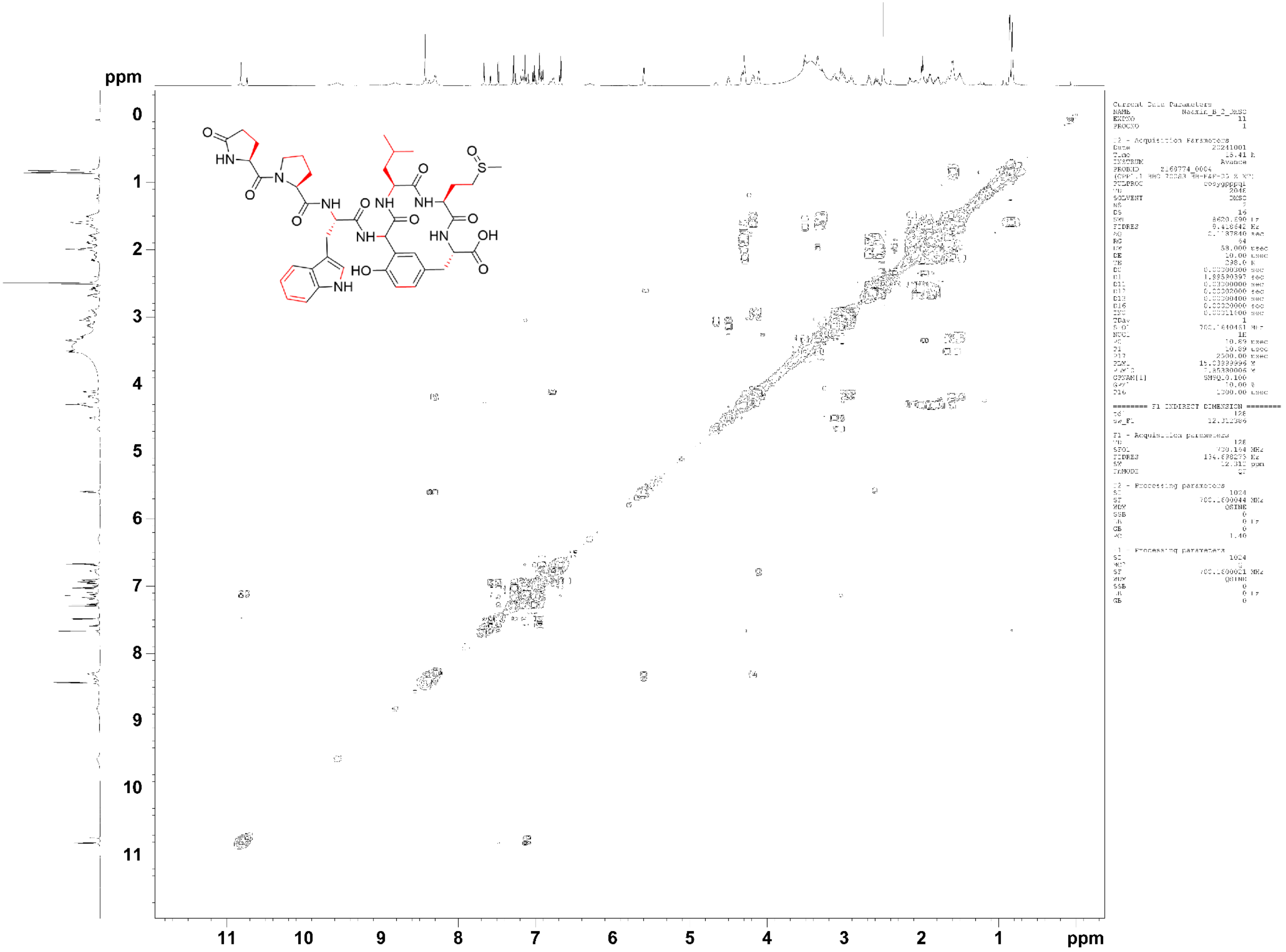


**Figure S4. COSY NMR spectrum of nanamin B (DMSO-$d_6$, 700 MHz, 298 K)**

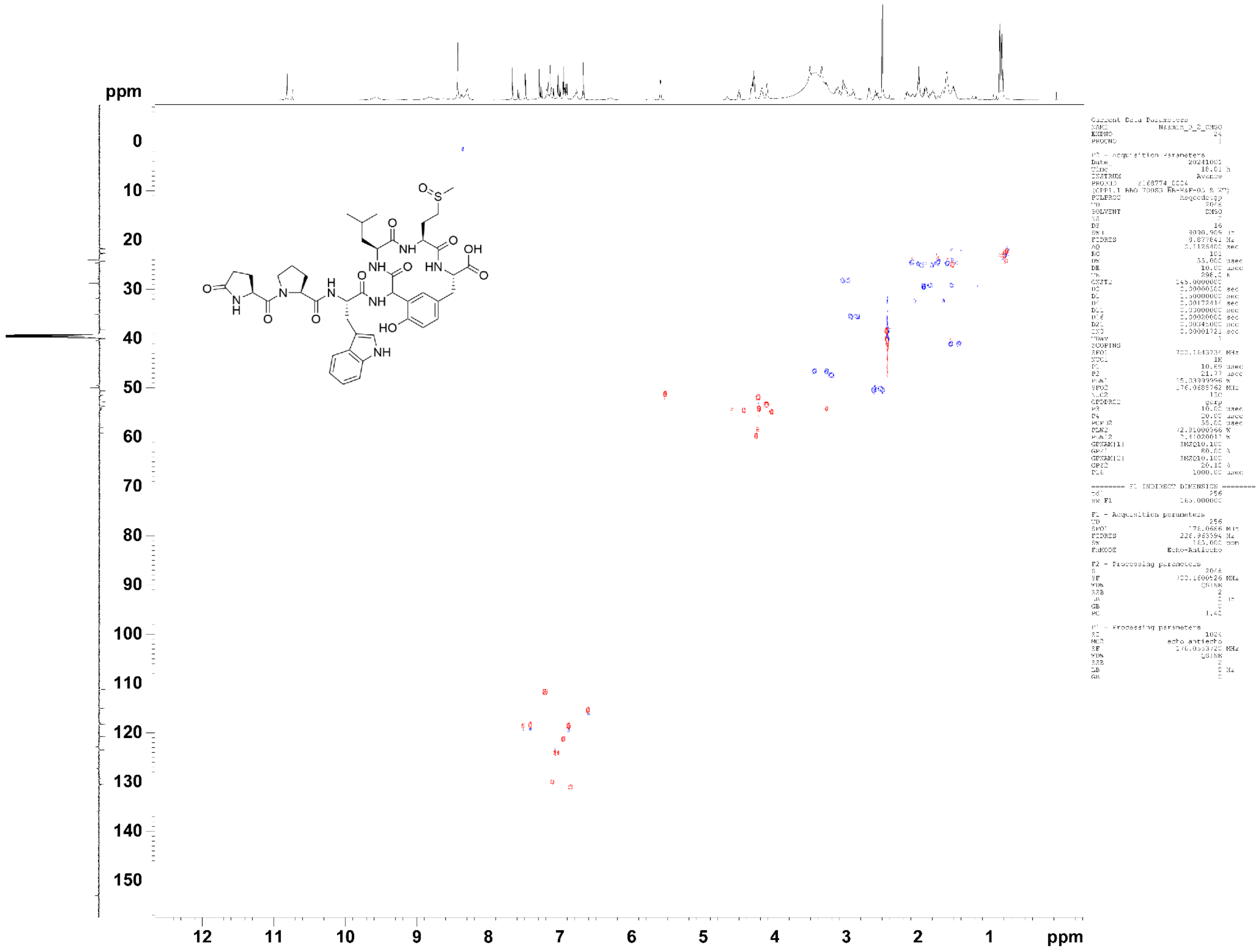


**Figure S5. HSQC-DEPT NMR spectrum of nanamin B (DMSO-$d_6$, 700 MHz, 298 K)**

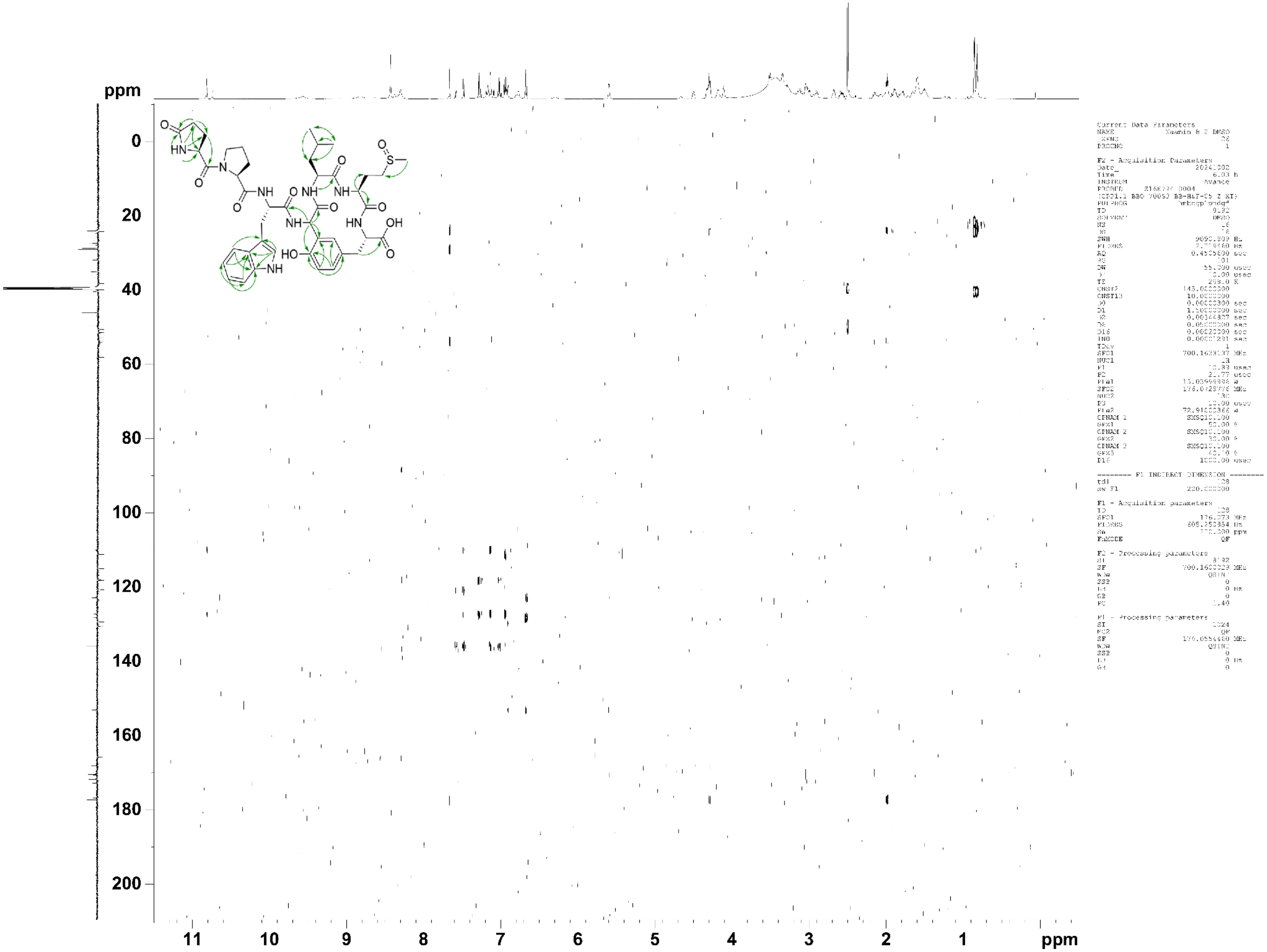


**Figure S6. HMBC NMR spectrum of nanamin B (DMSO-$d_6$, 700 MHz, 298 K)**

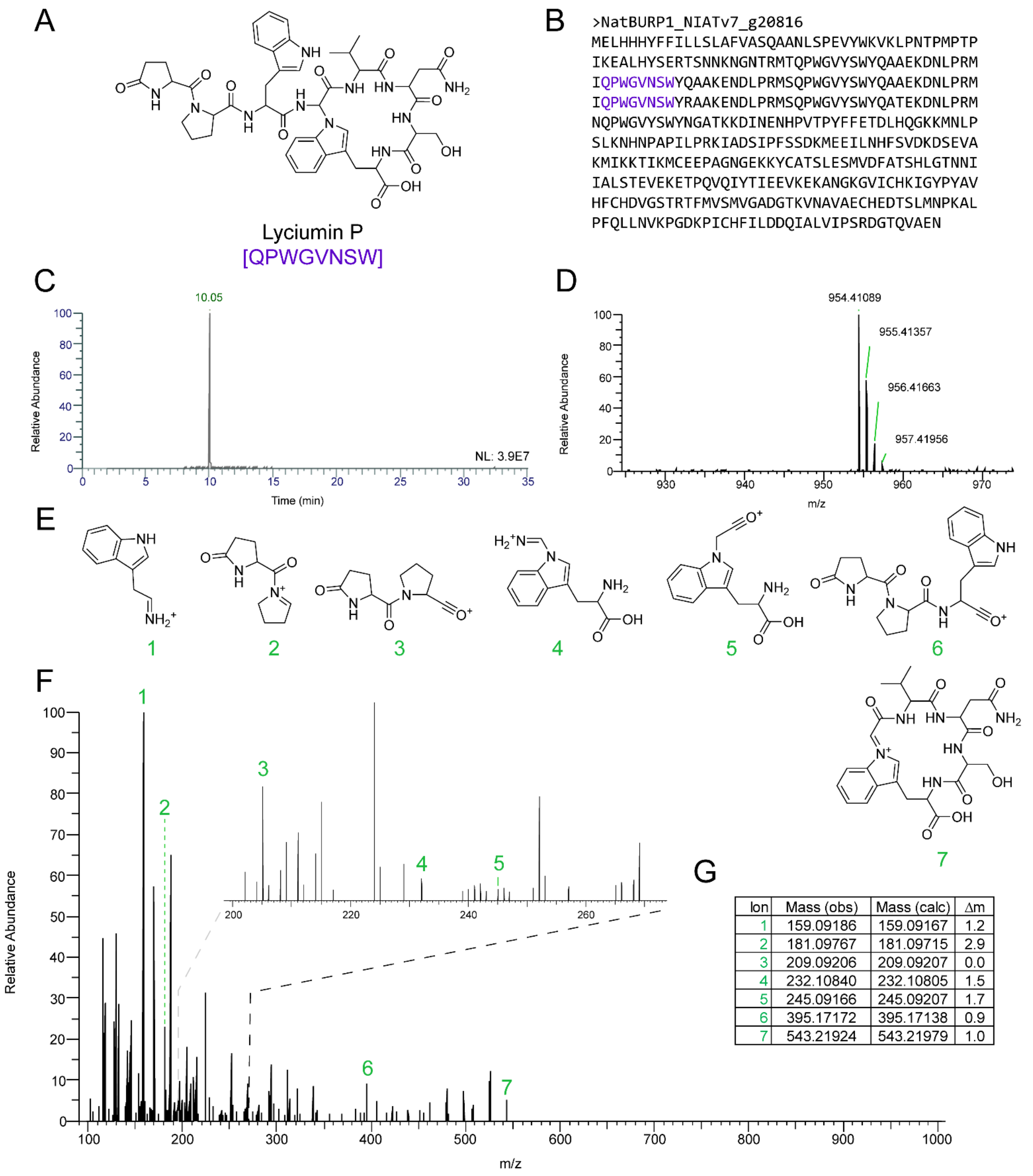


| Ion | Mass (obs) | Mass (calc) | Δm |
|---|---|---|---|
| 1 | 159.09186 | 159.09167 | 1.2 |
| 2 | 181.09767 | 181.09715 | 2.9 |
| 3 | 209.09206 | 209.09207 | 0.0 |
| 4 | 232.10840 | 232.10805 | 1.5 |
| 5 | 245.09166 | 245.09207 | 1.7 |
| 6 | 395.17172 | 395.17138 | 0.9 |
| 7 | 543.21924 | 543.21979 | 1.0 |

**Figure S7. MS/MS analysis of lyciumin P.** (**A**) Structure of Lyciumin P. (**B**) Sequence of NatBURP1. (**C**) EIC (954.41044 m/z, 5 ppm, +p ESI Full MS, [300-1500]) of peptide extracts of *N. attenuata* roots. (**D**) MS analysis of Lyciumin P at RT 10.1 min. (**E**) Predicted daughter ions. (**F**) MS/MS spectra. (**G**) Observed vs. calculated daughter ion masses.

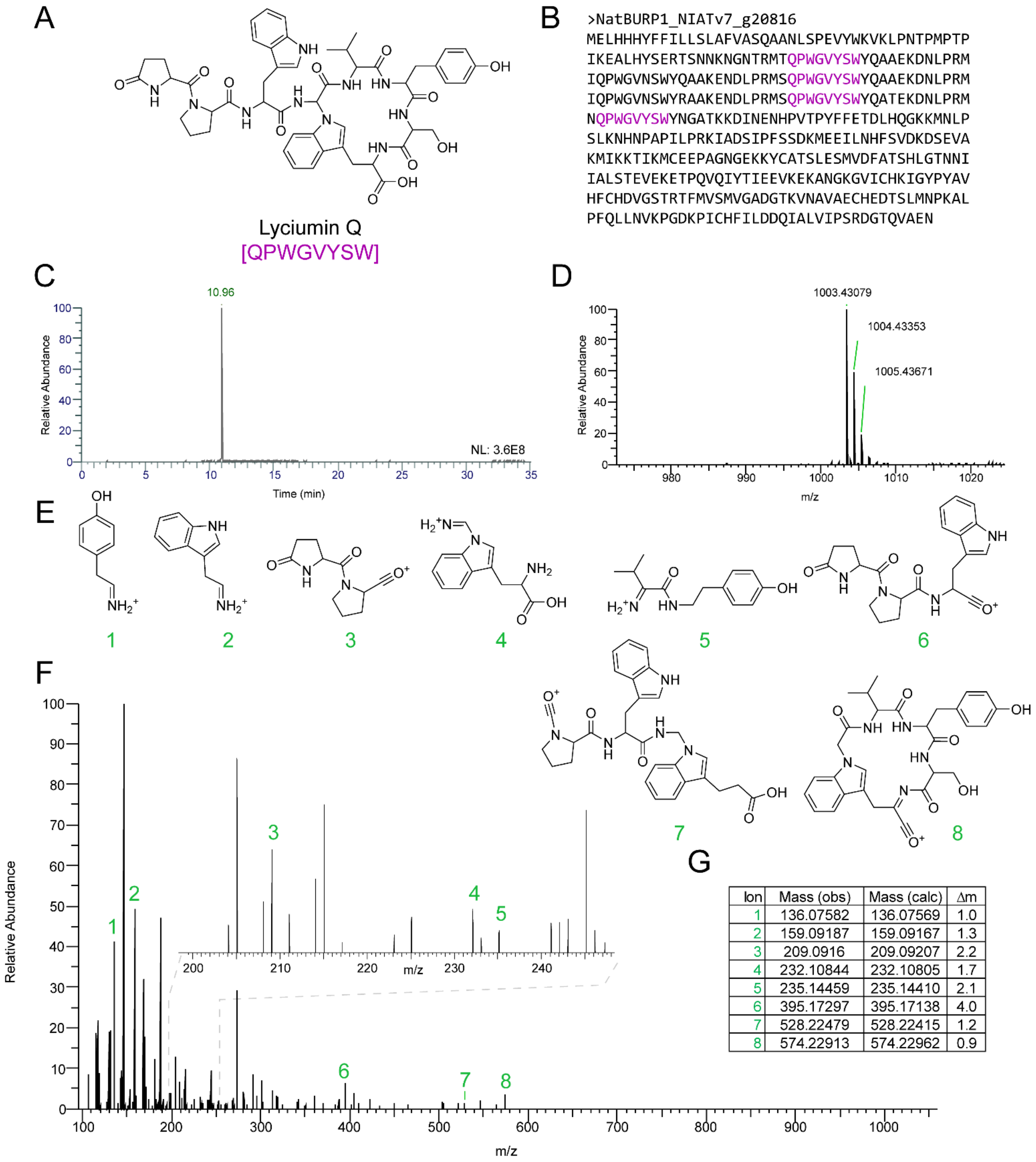


>NatBURP1_NIATv7_g20816
MELHHHYFFILLSLAFVASQAANLSPEVYWKVKLPNTPMPTP
IKEALHYSERTSNNKNGNTRMTQPWGVYSWYQAAEKDNLPRM
IQPWGVNSWYQAAKENDLPRMSQPWGVYSWYQAAEKDNLPRM
IQPWGVNSWYRAAKENDLPRMSQPWGVYSWYQATEKDNLPRM
NQPWGVYSWYNGATKKDINENHPVTPYFFETDLHQGKKMNLP
SLKNHNPAPILPRKIADSIPFSSDKMEEILNHFSVDKDSEVA
KMIKKTIKMCEEPAGNGEKKYCATSLESMVDFATSHLGTNNI
IALSTEVEKETPQVQIYTIEEVKEKANGKGVICHKIGYPYAV
HFCHDVGSTRTFMVSMVGADGTKVNAVAECHEDTSLMNPKAL
PFQLLNVKPGDKPICHFILDDQIALVIPSRDGTQVAEN

| Ion | Mass (obs) | Mass (calc) | Δm |
|---|---|---|---|
| 1 | 136.07582 | 136.07569 | 1.0 |
| 2 | 159.09187 | 159.09167 | 1.3 |
| 3 | 209.0916 | 209.09207 | 2.2 |
| 4 | 232.10844 | 232.10805 | 1.7 |
| 5 | 235.14459 | 235.14410 | 2.1 |
| 6 | 395.17297 | 395.17138 | 4.0 |
| 7 | 528.22479 | 528.22415 | 1.2 |
| 8 | 574.22913 | 574.22962 | 0.9 |

**Figure S8. MS/MS analysis of lyciumin Q.** (**A**) Structure of lyciumin Q. (**B**) Sequence of NatBURP1. (**C**) EIC (1003.43084 m/z, 5 ppm, +p ESI Full MS, [300-1500]) of peptide extracts of *N. attenuata* roots. (**D**) MS analysis of lyciumin Q at RT 11.0 min. (**E**) Predicted daughter ions. (**F**) MS/MS spectra. (**G**) Observed vs. calculated daughter ion masses.

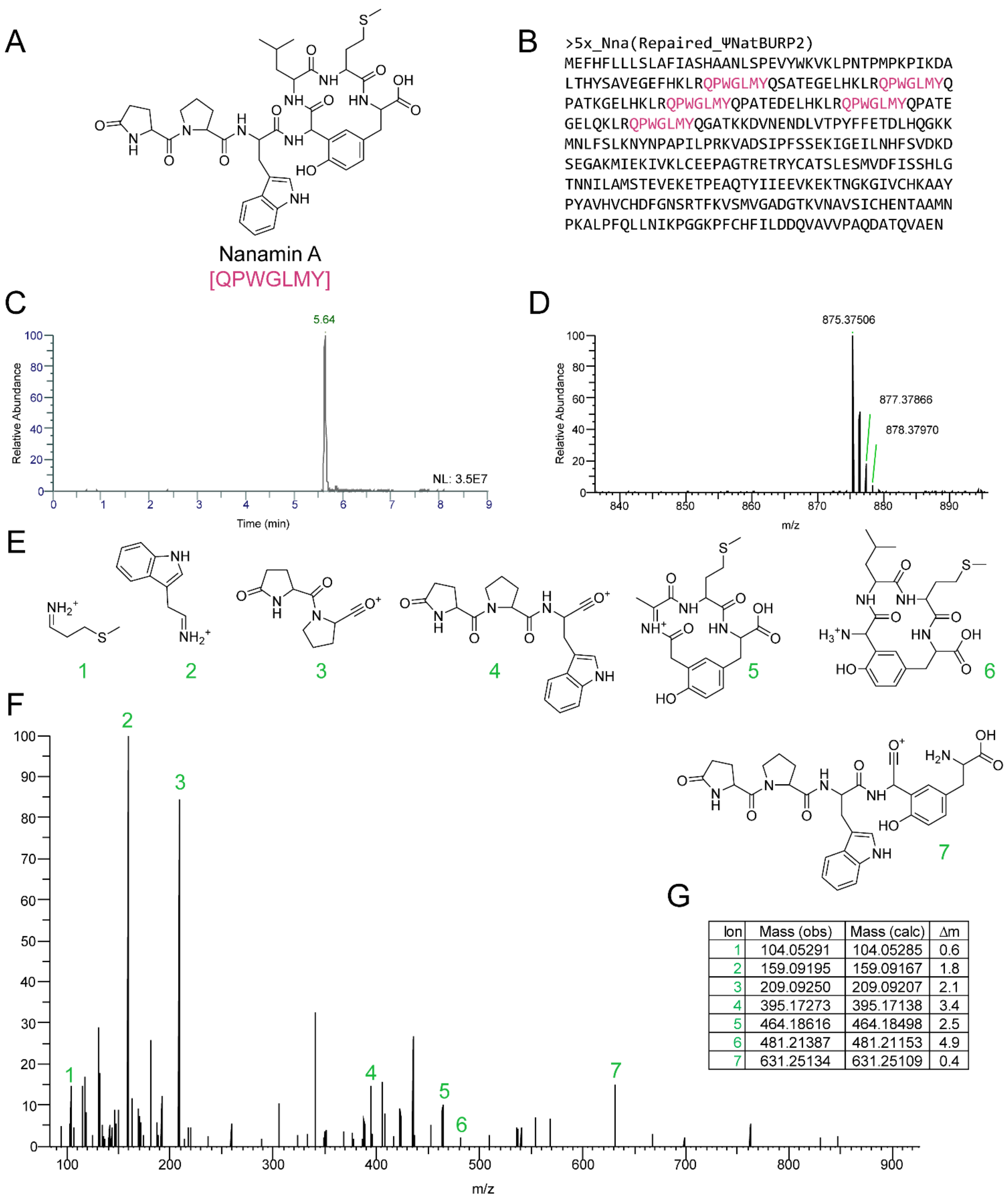


| Ion | Mass (obs) | Mass (calc) | Δm |
|---|---|---|---|
| 1 | 104.05291 | 104.05285 | 0.6 |
| 2 | 159.09195 | 159.09167 | 1.8 |
| 3 | 209.09250 | 209.09207 | 2.1 |
| 4 | 395.17273 | 395.17138 | 3.4 |
| 5 | 464.18616 | 464.18498 | 2.5 |
| 6 | 481.21387 | 481.21153 | 4.9 |
| 7 | 631.25134 | 631.25109 | 0.4 |

**Figure S9. MS/MS analysis of nanamin A.** (**A**) Structure of nanamin A. (**B**) Sequence of 5x_Nna. (**C**) EIC (875.37564 m/z, 5 ppm, +p ESI Full MS, [300-1500]) of *N. benthamiana* agroinfiltrated with 5x_Nna. (**D**) MS analysis of nanamin A at RT 5.6 min. (**E**) Predicted daughter ions. (**F**) MS/MS spectra. (**G**) Observed vs. calculated daughter ion masses.

A

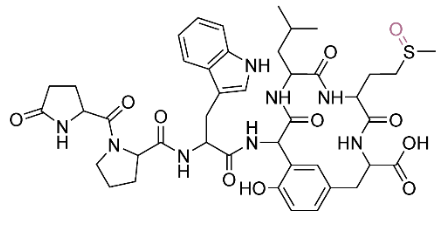


Nanamin B
[QPWGL(MetO)Y]

B

```
>NclBURP2_(N.clevelandii_TW30)
MEFHFLLLSLAFIASHAANLSPEVYWKVKLPNTPMPKPIKDALTHYSA
VEGEFPKLRQPWGLMYQPATEGELQKLRQPWGLMYQPATEGELQKLRQ
PWGLMYQPATEDELHKLRQPWGLMYKPATEGELHKLRQPWGLMYQPAT
EGELHKLRQPWGLMYQPATEDELHKLRQPWGLMYKPATEGELHKLRQP
WGLMYQPATEGELHKLRQPWGLMYQPATEGELQKLRQPWGLMYQPATE
GELQKLRQPWGLMYQPATEDELHKLRQPWGLMYKPATEGELHKLRQPW
GLMYQPATEGELQKLRQPWGLMYQPATEGELHKLRQPWGLMYQPATEN
ELHKLRQPWGLMYQPATEGELHKLRQPWGLMYQGATKKDVNENGLVTP
YFFETDLHQGKKMNLLSLKNYNPAPILPRKVADSIPFSSEKIGEILNQ
FSVDKDSEGAKMIEKIVKLCEEPAGTRETRYCATSLESMVDFISSHLG
TNNILAMSTEVEKETPEAQTYIIEEVKEKTNGKGIVCHKAAYPYAVHV
CHDFGNSRTFKVSMVGADGTKVNAVSICHENTAAMNPKALPFQLLNIK
PGGKPFCHFILDDQVAIVPAQDATQVAEN
```

C

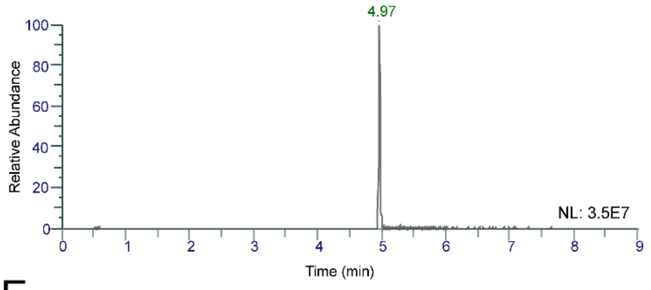


D

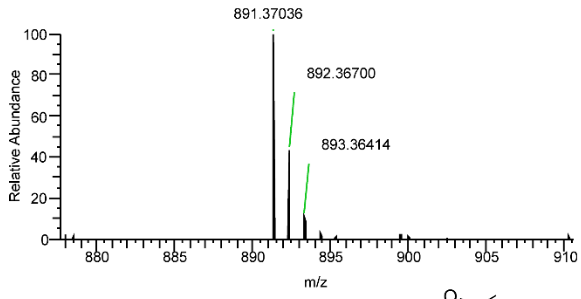


E

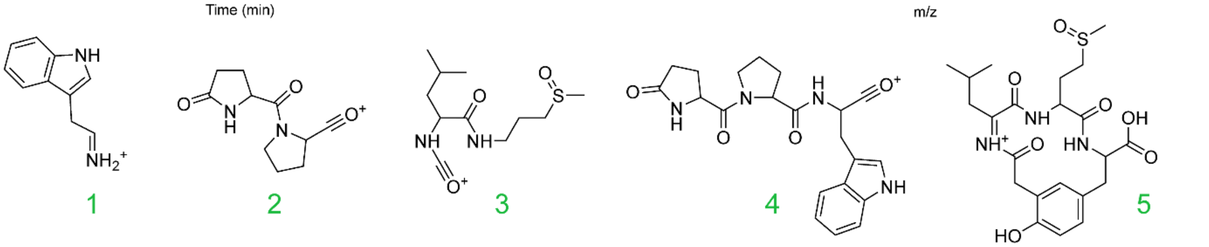


F

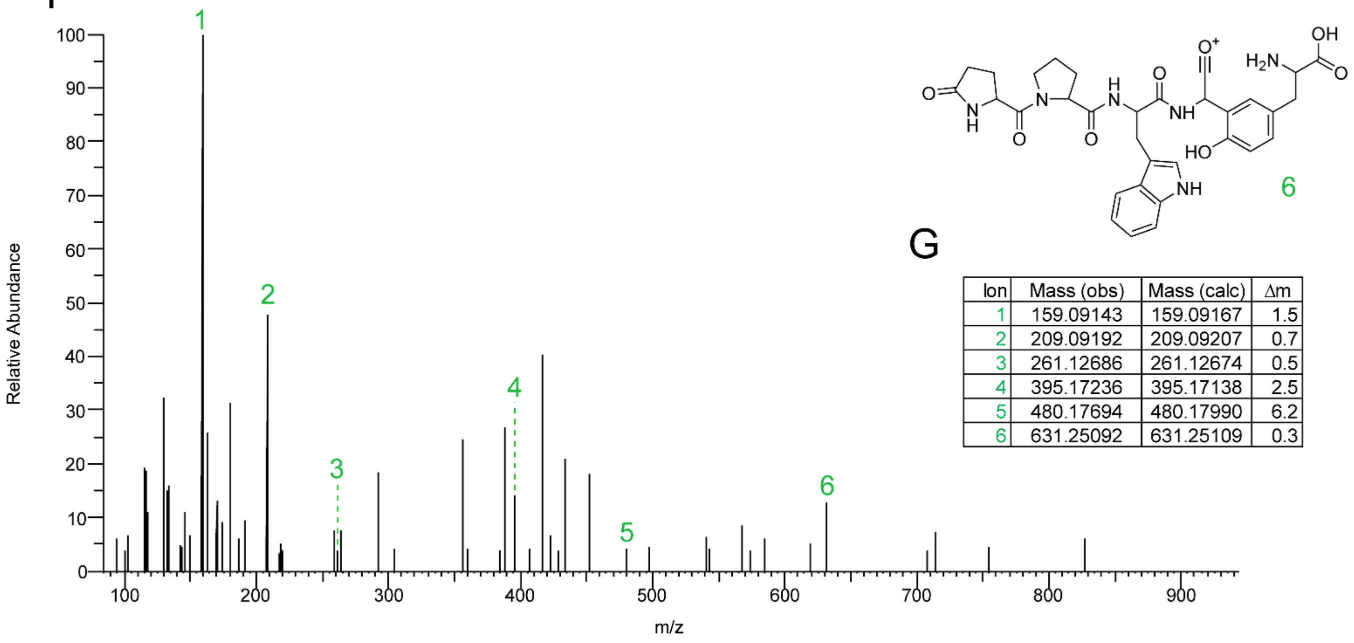


G

| Ion | Mass (obs) | Mass (calc) | Δm |
|---|---|---|---|
| 1 | 159.09143 | 159.09167 | 1.5 |
| 2 | 209.09192 | 209.09207 | 0.7 |
| 3 | 261.12686 | 261.12674 | 0.5 |
| 4 | 395.17236 | 395.17138 | 2.5 |
| 5 | 480.17694 | 480.17990 | 6.2 |
| 6 | 631.25092 | 631.25109 | 0.3 |

**Figure S10. MS/MS analysis of nanamin B.** (**A**) Structure of nanamin A. (**B**) Sequence of NclBURP2. (**C**) EIC (890.36328 m/z, 5 ppm, +p ESI Full MS, [300-1500]) of *N. benthamiana* agroinfiltrated with NclBURP2. (**D**) MS analysis of nanamin B at RT 5.0 min. (**E**) Predicted daughter ions. (**F**) MS/MS spectra. (**G**) Observed vs. calculated daughter ion masses.

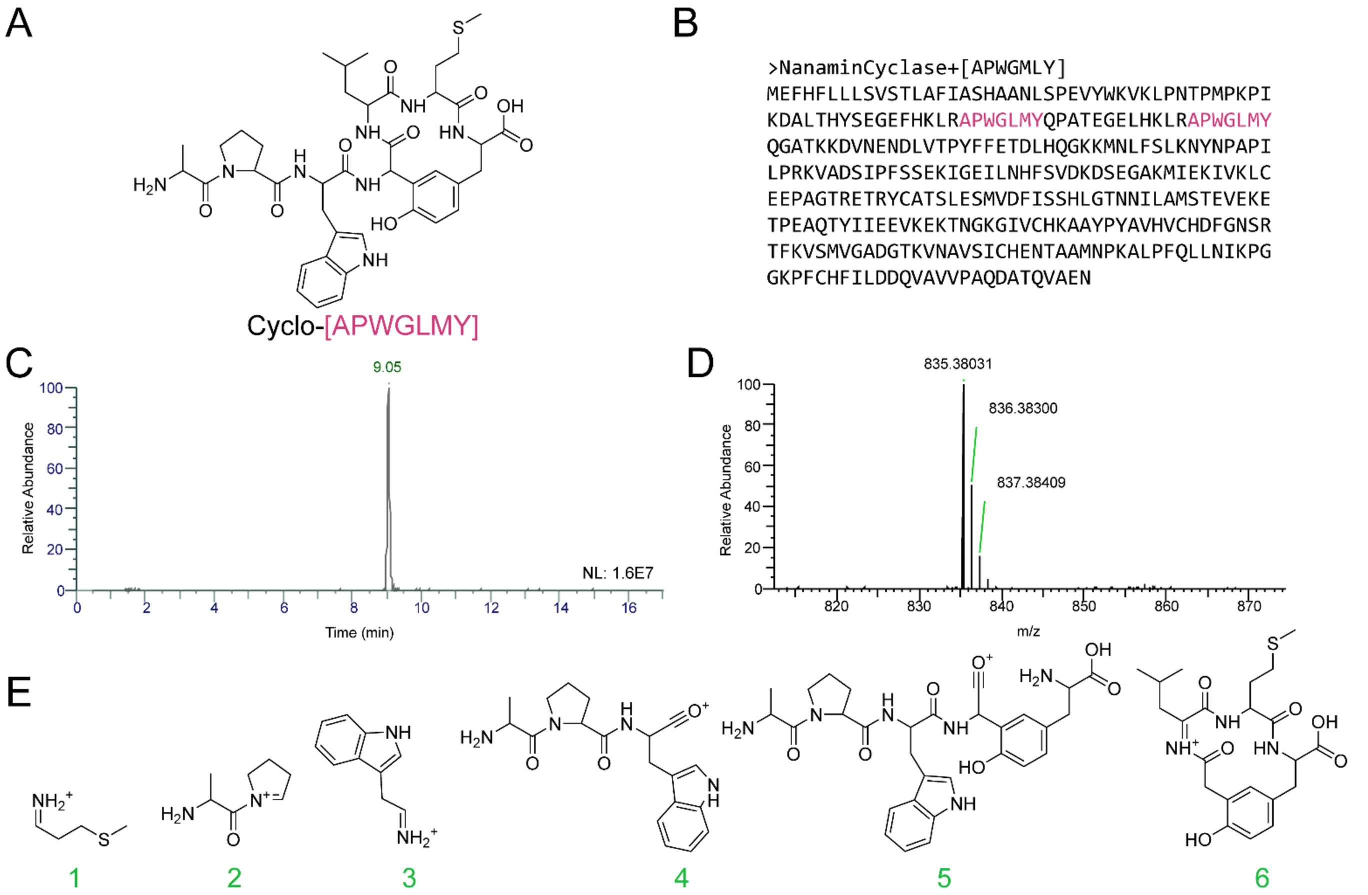


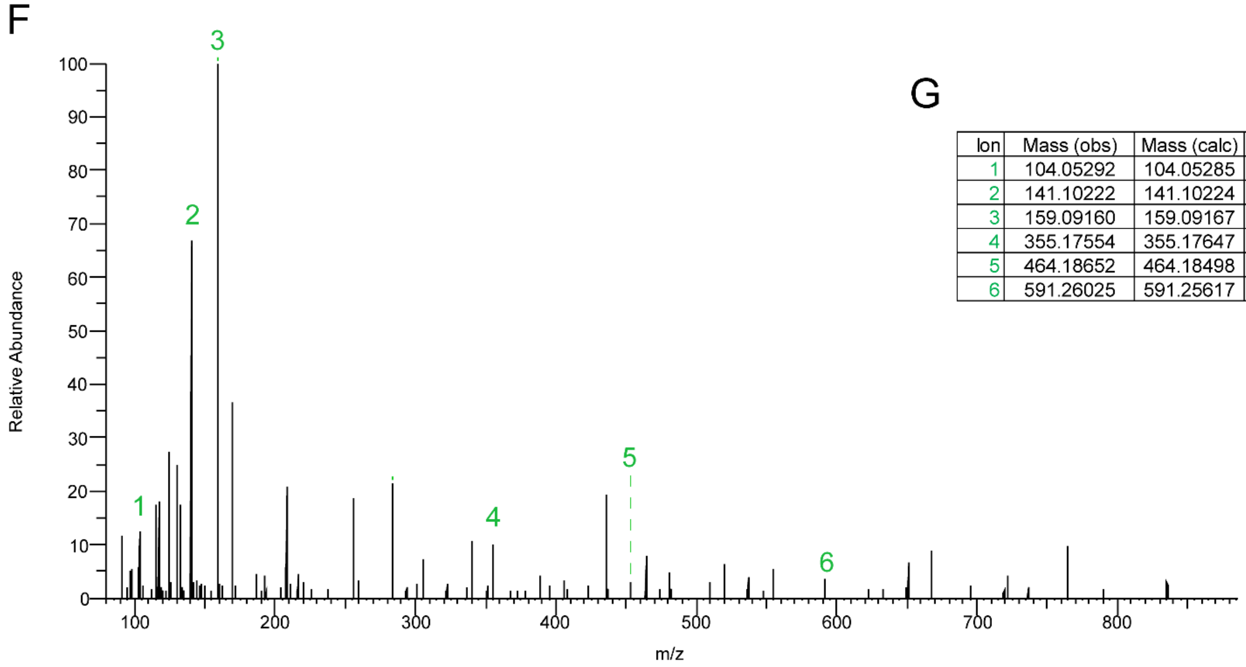


G

| Ion | Mass (obs) | Mass (calc) | Δm |
|---|---|---|---|
| 1 | 104.05292 | 104.05285 | 0.7 |
| 2 | 141.10222 | 141.10224 | 0.1 |
| 3 | 159.09160 | 159.09167 | 0.4 |
| 4 | 355.17554 | 355.17647 | 2.6 |
| 5 | 464.18652 | 464.18498 | 3.3 |
| 6 | 591.26025 | 591.25617 | 6.9 |

**Figure S11. MS/MS analysis of cyclo-[APWGLMY].** (**A**) Putative structure of cyclo-[AWPGLMY]. (**B**) Sequence of NanaminCyclase+[APWGLMY]. (**C**) EIC (418.19400 m/z, 5 ppm, +p ESI Full MS, [300-1500]) of *N. benthamiana* agroinfiltrated with NanaminCyclase+[APWGLMY]. (**D**) MS analysis of cyclo-[APWGLMY] at RT 9.1 min. (**E**) Predicted daughter ions. (**F**) MS/MS spectra. (**G**) Observed vs. calculated daughter ion masses.

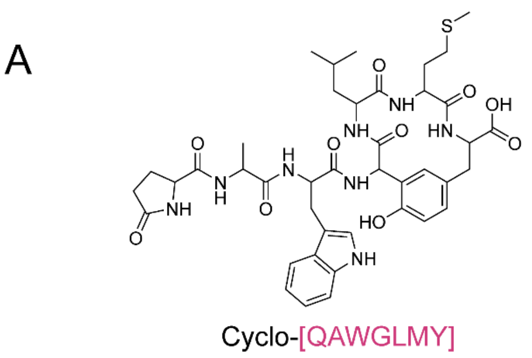


B

>NanaminCyclase+[QAWGLMY]
MEFHFLLLSVSTLAFIASHAANLSPEVYWKVKLPNTPMPKPI
KDALTHYSEGEFHKLRQAWGLMYQPATEGELHKLRQAWGLMY
QGATKKDVNENDLVTPYFFETDLHQGKKMNLFSLKNYNPAPI
LPRKVADSIPFSSEKIGEILNHFSVDKDSEGAKMIEKIVKLC
EEPAGTRETRYCATSLESMVDFISSHLGTNNILAMSTEVEKE
TPEAQTYIIEEVKEKTNGKGIVCHKAAYPYAVHVCHDFGNSR
TFKVSMVGADGTKVNAVSICHENTAAMNPKALPFQLLNIKPG
GKPFCHFILDDQVAVVPAQDATQVAEN

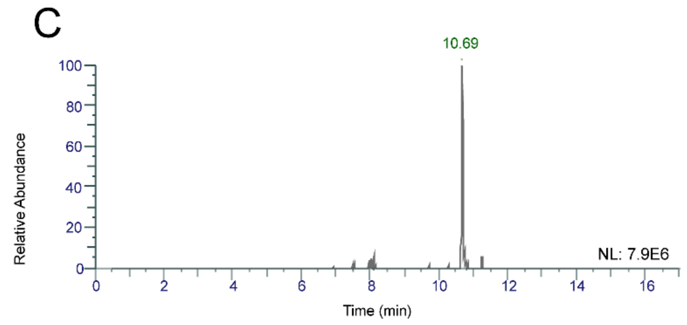


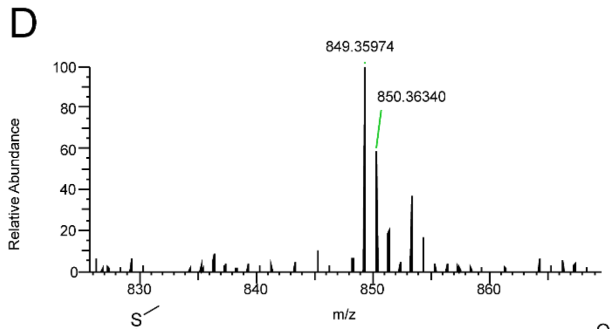


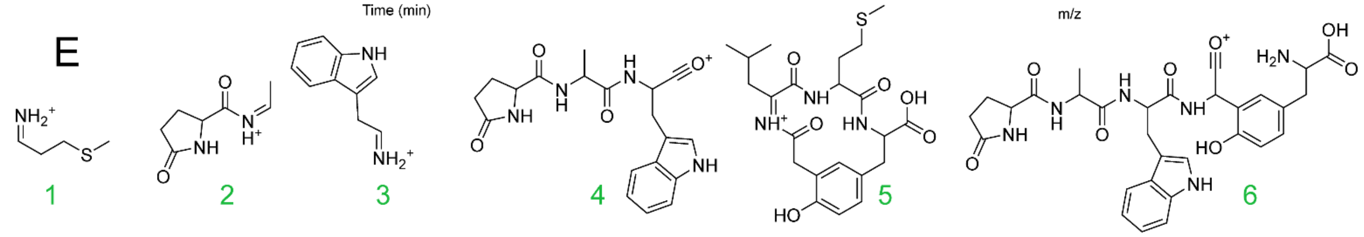


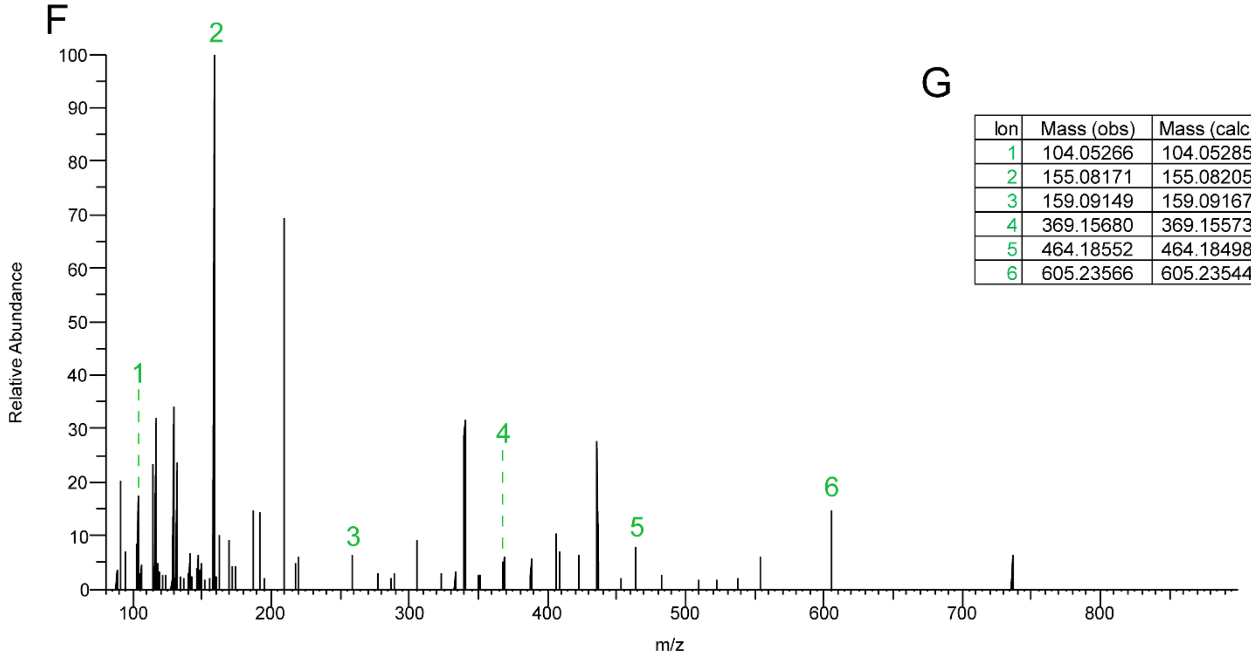


G

| Ion | Mass (obs) | Mass (calc) | Δm |
|---|---|---|---|
| 1 | 104.05266 | 104.05285 | 1.8 |
| 2 | 155.08171 | 155.08205 | 2.2 |
| 3 | 159.09149 | 159.09167 | 1.1 |
| 4 | 369.15680 | 369.15573 | 2.9 |
| 5 | 464.18552 | 464.18498 | 1.2 |
| 6 | 605.23566 | 605.23544 | 0.4 |

**Figure S12. MS/MS analysis of cyclo-[QAWGLMY].** (**A**) Putative structure of cyclo-[QAWGLMY]. (**B**) Sequence of NanaminCyclase+[QAWGLMY]. (**C**) EIC (849.35999 m/z, 5 ppm, +p ESI Full MS, [300-1500]) of *N. benthamiana* agroinfiltrated with NanaminCyclase+[QAWGLMY]. (**D**) MS analysis of cyclo-[QAWGLMY] at RT 10.7 min. (**E**) Predicted daughter ions. (**F**) MS/MS spectra. (**G**) Observed vs. calculated daughter ion masses.

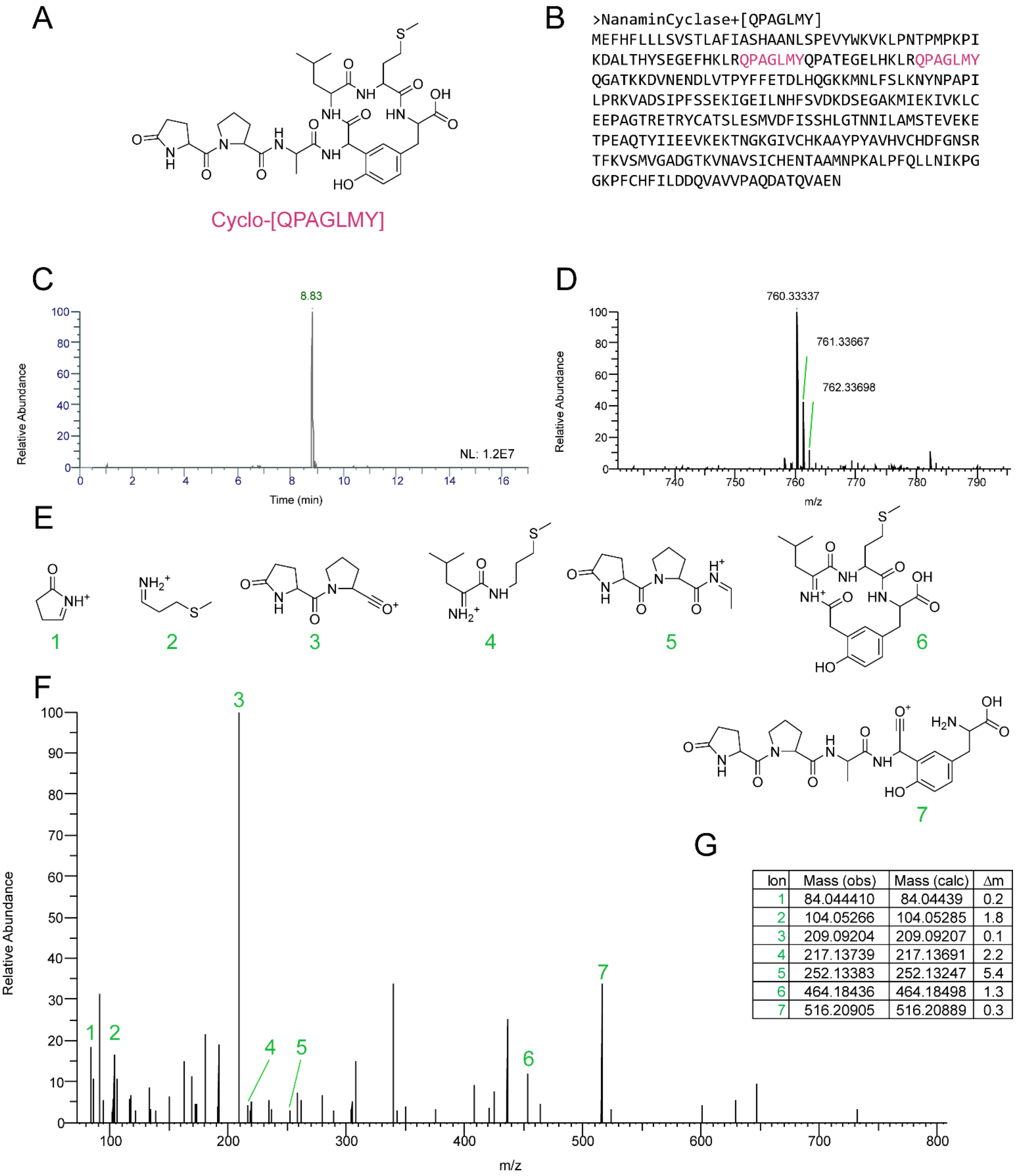


| Ion | Mass (obs) | Mass (calc) | Δm |
|---|---|---|---|
| 1 | 84.044410 | 84.04439 | 0.2 |
| 2 | 104.05266 | 104.05285 | 1.8 |
| 3 | 209.09204 | 209.09207 | 0.1 |
| 4 | 217.13739 | 217.13691 | 2.2 |
| 5 | 252.13383 | 252.13247 | 5.4 |
| 6 | 464.18436 | 464.18498 | 1.3 |
| 7 | 516.20905 | 516.20889 | 0.3 |

**Figure S13. MS/MS analysis of cyclo-[QPAGLMY].** (**A**) Putative structure of cyclo-[QPAGLMY]. (**B**) Sequence of NanaminCyclase+[QPAGLMY]. (**C**) EIC (760.33344 m/z, 5 ppm, +p ESI Full MS, [300-1500]) of *N. benthamiana* agroinfiltrated with NanaminCyclase+[QPAGLMY]. (**D**) MS analysis of cyclo-[QPAGLMY] at RT 8.8 min. (**E**) Predicted daughter ions. (**F**) MS/MS spectra. (**G**) Observed vs. calculated daughter ion masses.

A

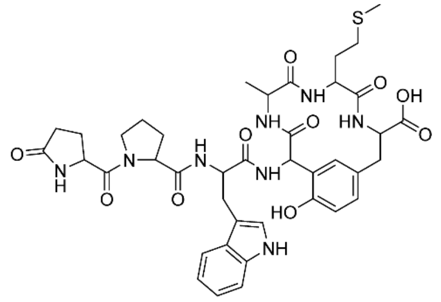


Cyclo-[QPWGAMY]

B

```
>NanaminCyclase+[QPWGAMY]
MEFHFLLLSVSTLAFIASHAANLSPEVYWKVKLPNTPMPKPI
KDALTHYSEGEFHKLRQPWGAMYQPATEGELHKLRQPWGAMY
QGATKKDVNENDLVTPYFFETDLHQGKKMNLFSLKNYNPAPI
LPRKVADSIPFSSEKIGEILNHFSVDKDSEGAKMIEKIVKLC
EEPAGTRETRYCATSLESMVDFISSHLGTNNILAMSTEVEKE
TPEAQTYIIEEVKEKTNGKGIVCHKAAYPYAVHVCHDFGNSR
TFKVSMVGADGTKVNAVSICHENTAAMNPKALPFQLLNIKPG
GKPFCHFILDDQVAVVPAQDATQVAEN
```

C

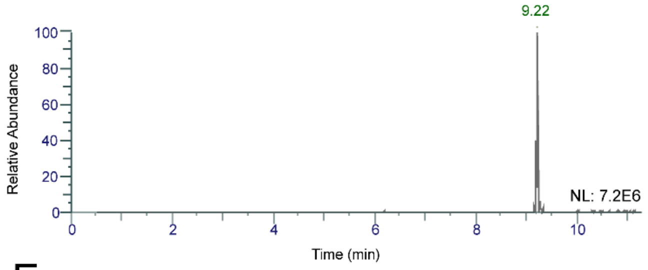


D

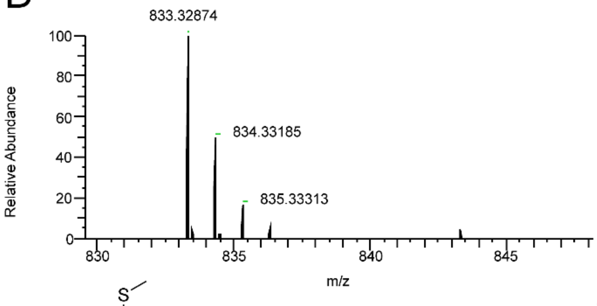


E

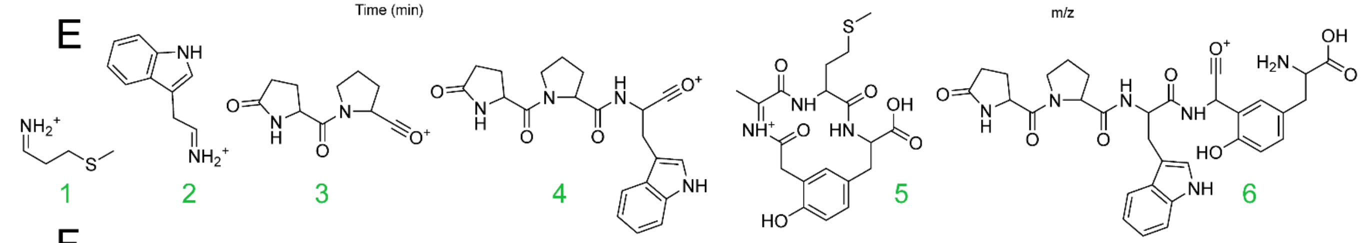


F

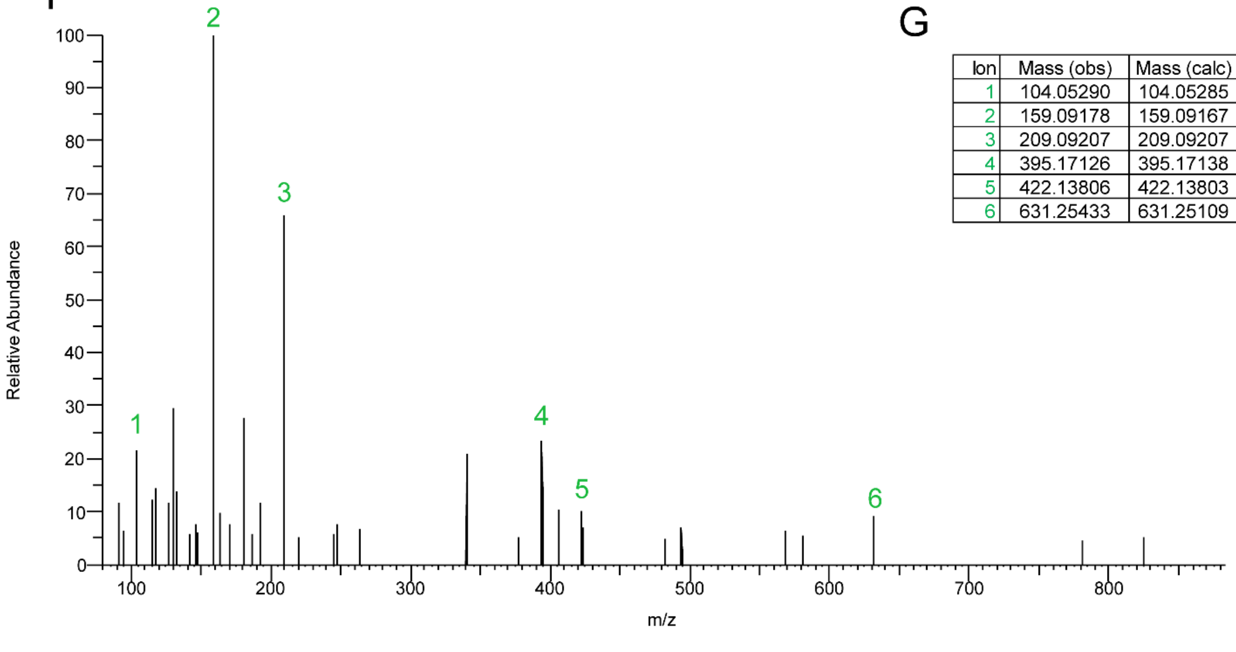


G

| Ion | Mass (obs) | Mass (calc) | Δm |
|---|---|---|---|
| 1 | 104.05290 | 104.05285 | 0.5 |
| 2 | 159.09178 | 159.09167 | 0.7 |
| 3 | 209.09207 | 209.09207 | 0.0 |
| 4 | 395.17126 | 395.17138 | 0.3 |
| 5 | 422.13806 | 422.13803 | 0.1 |
| 6 | 631.25433 | 631.25109 | 5.1 |

**Figure S14. MS/MS analysis of cyclo-[QPWGAMY].** (**A**) Putative structure of cyclo-[QPWGAMY]. (**B**) Sequence of NanaminCyclase+[QPWGAMY]. (**C**) EIC (833.32869 m/z, 5 ppm, +p ESI Full MS, [300-1500]) of *N. benthamiana* agroinfiltrated with NanaminCyclase+[QPWGAMY]. (**D**) MS analysis of cyclo-[QPWGAMY] at RT 9.2 min. (**E**) Predicted daughter ions. (**F**) MS/MS spectra. (**G**) Observed vs. calculated daughter ion masses.

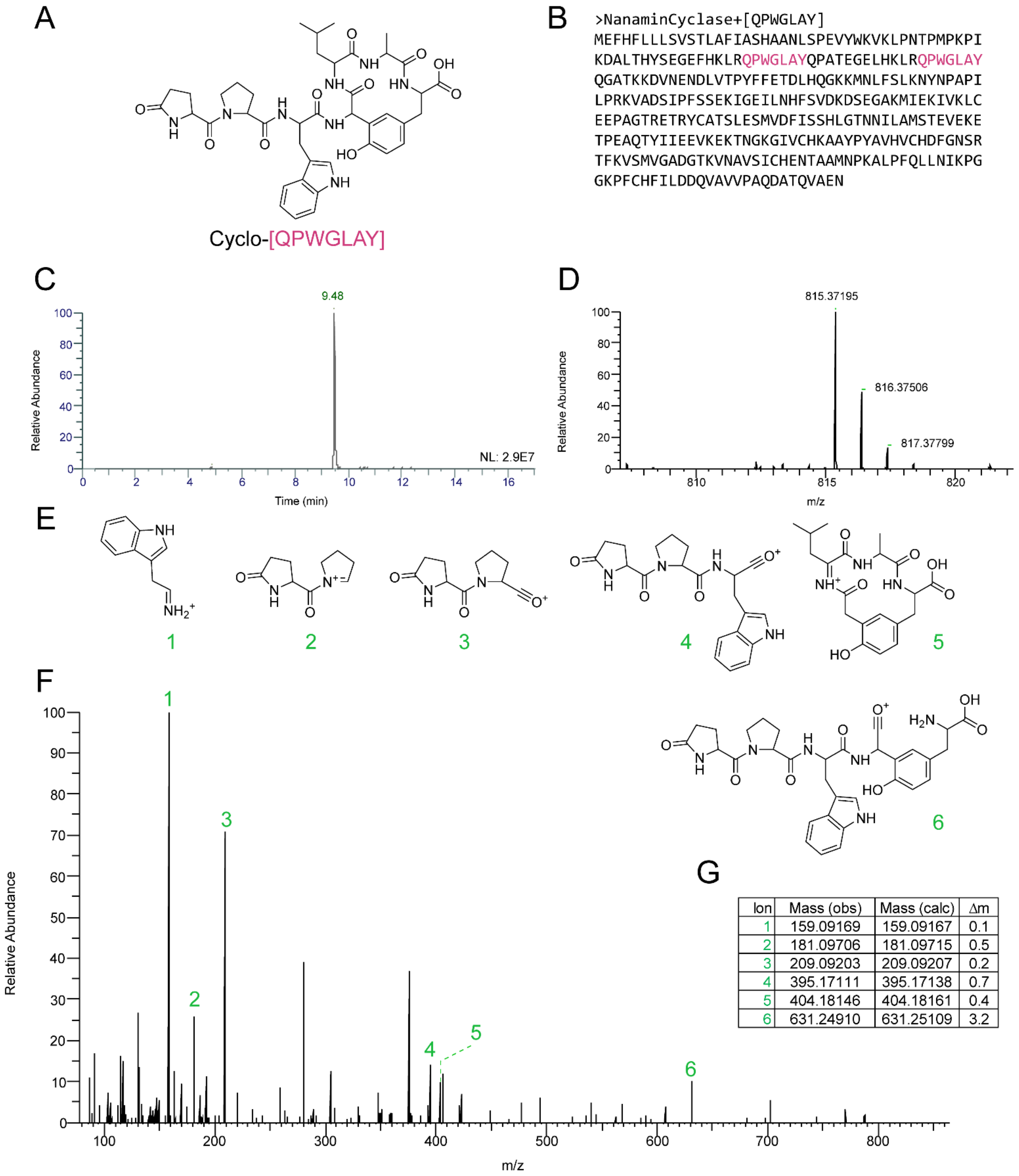


| Ion | Mass (obs) | Mass (calc) | Δm |
|---|---|---|---|
| 1 | 159.09169 | 159.09167 | 0.1 |
| 2 | 181.09706 | 181.09715 | 0.5 |
| 3 | 209.09203 | 209.09207 | 0.2 |
| 4 | 395.17111 | 395.17138 | 0.7 |
| 5 | 404.18146 | 404.18161 | 0.4 |
| 6 | 631.24910 | 631.25109 | 3.2 |

**Figure S15. MS/MS analysis of cyclo-[QPWGLAY].** (**A**) Putative structure of cyclo-[QPWGLAY]. (**B**) Sequence of NanaminCyclase+[QPWGLAY]. (**C**) EIC (815.37227 m/z, 5 ppm, +p ESI Full MS, [300-1500]) of *N. benthamiana* agroinfiltrated with NanaminCyclase+[QPWGLAY]. (**D**) MS analysis of cyclo-[QPWGLAY] at RT 9.5 min. (**E**) Predicted daughter ions. (**F**) MS/MS spectra. (**G**) Observed vs. calculated daughter ion masses.

A

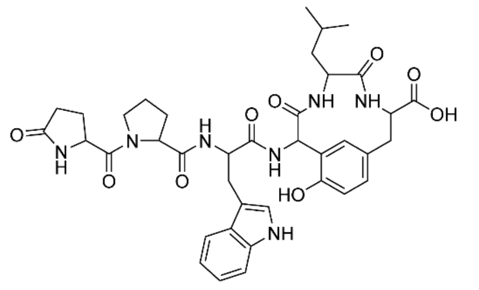


B

```
>NanaminCyclase+[QPGLMY]
MEFHFLLLSLAFIASHAANLSPEVYWKVKLPNTPMPKPIKDA
LTHYSAVEGEFHKLRQPGLMYQSATEGELHKLRQPGLMYQPT
TEGELHKLRQPGLMYQPATEGEIHKLRQPGLMYQLATEGELR
KLRQPGLMYQGATKKDVNENDLVTPYFFETDLHQGKKMNLFS
LKNYNPAPILPRKVADSIPFSSEKIGEILNHFSVDKDSEGAK
MIEKIVKLCEEPAGTRETRYCATSLESMVDFISSHLGTNNIL
AMSTEVEKETPEAQTYIIEEVKEKTNGKGIVCHKAAYPYAVH
VCHDFGNSRTFKVSMVGADGTKVNAVSICHENTAAMNPKALP
FQLLNIKPGGKPFCHFILDDQVAVVPAQDATQVAEN
```

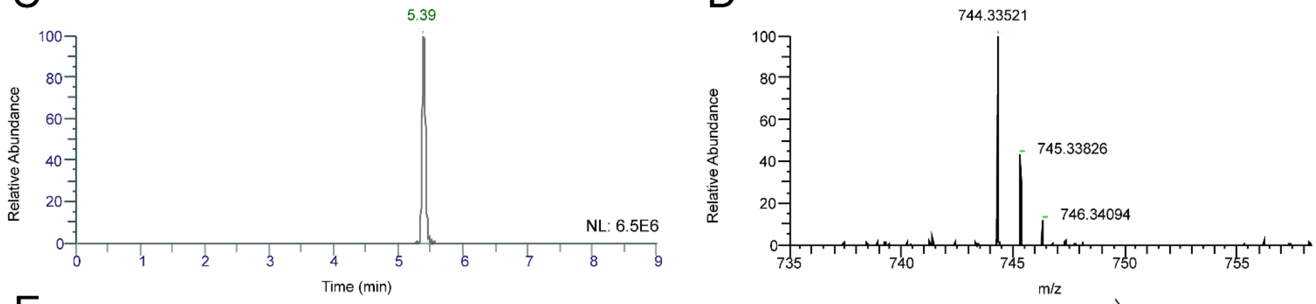


E

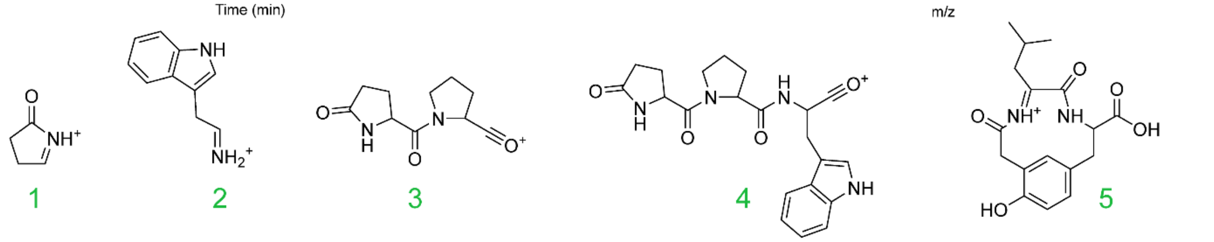


F

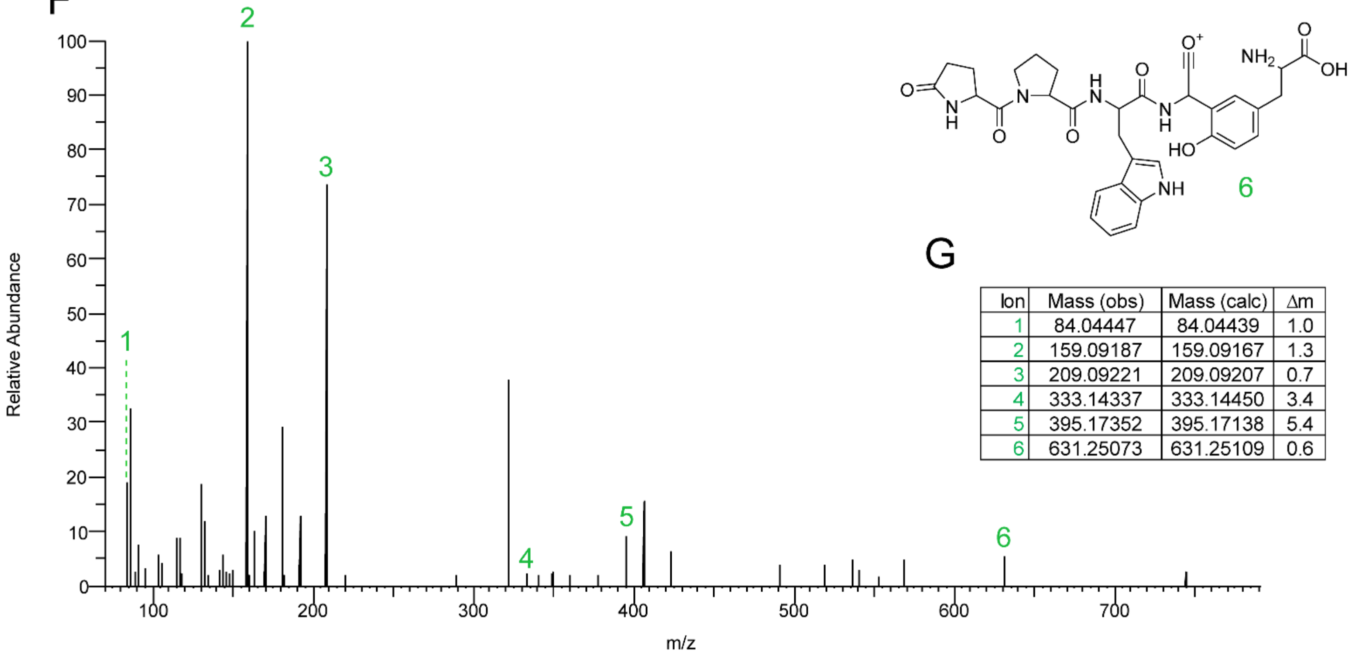


G

| Ion | Mass (obs) | Mass (calc) | Δm |
|---|---|---|---|
| 1 | 84.04447 | 84.04439 | 1.0 |
| 2 | 159.09187 | 159.09167 | 1.3 |
| 3 | 209.09221 | 209.09207 | 0.7 |
| 4 | 333.14337 | 333.14450 | 3.4 |
| 5 | 395.17352 | 395.17138 | 5.4 |
| 6 | 631.25073 | 631.25109 | 0.6 |

**Figure S16. MS/MS analysis of cyclo-[QPWGLY].** (**A**) Putative structure of cyclo-[QPWGLY]. (**B**) Sequence of NanaminCyclase+[QPWGLY]. (**C**) EIC (744.33515 m/z, 5 ppm, +p ESI Full MS, [300-1500]) of *N. benthamiana* agroinfiltrated with NanaminCyclase+[QPWGLY]. (D) MS analysis of cyclo-[QPWGLY] at RT 5.4 min. (**E**) Predicted daughter ions. (F) MS/MS spectra. (**G**) Observed vs. calculated daughter ion masses.

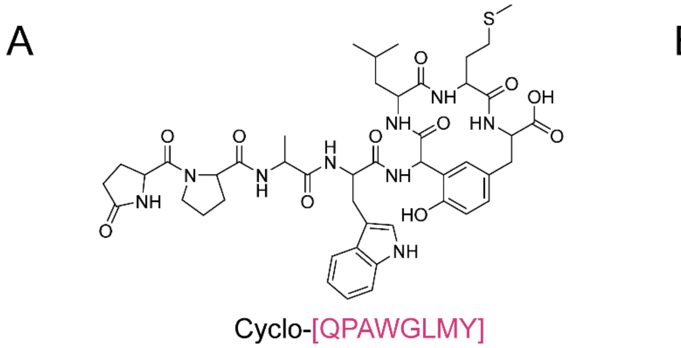


B

```
>NanaminCyclase+[QPAWGLMY]
MEFHFLLLSLAFIASHAANLSPEVYWKVKLPNTPMPKPIKDA
LTHYSAVEGEFHKLRQPAWGLMYQSATEGELHKLRQPAWGLM
YQPTTEGELHKLRQPAWGLMYQPATEGEIHKLRQPAWGLMYQ
LATEGELRKLRQPAWGLMYQGATKKDVNENDLVTPYFFETDL
HQGKKMNLFSLKNYNPAPILPRKVADSIPFSSEKIGEILNHF
SVDKDSEGAKMIEKIVKLCEEPAGTRETRYCATSLESMVDFI
SSHLGTNNILAMSTEVEKETPEAQTYIIEEVKEKTNGKGIVC
HKAAYPYAVHVCHDFGNSRTFKVSMVGADGTKVNAVSICHEN
TAAMNPKALPFQLLNIKPGGKPFCHFILDDQVAVVPAQDATQ
VAEN
```

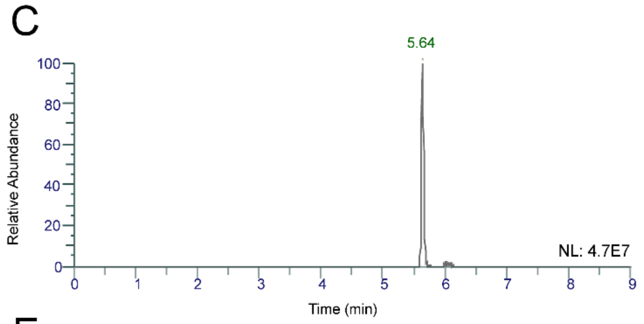


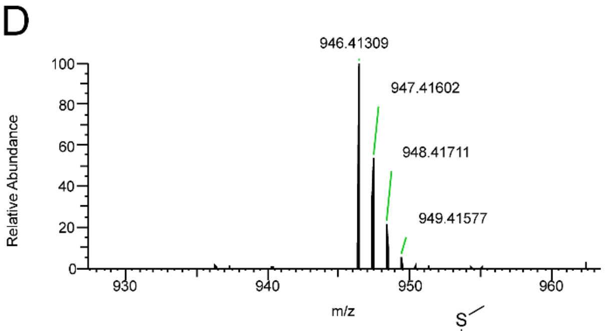


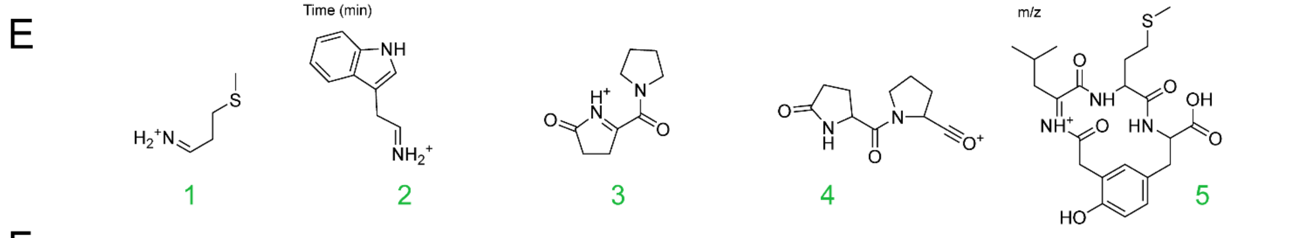


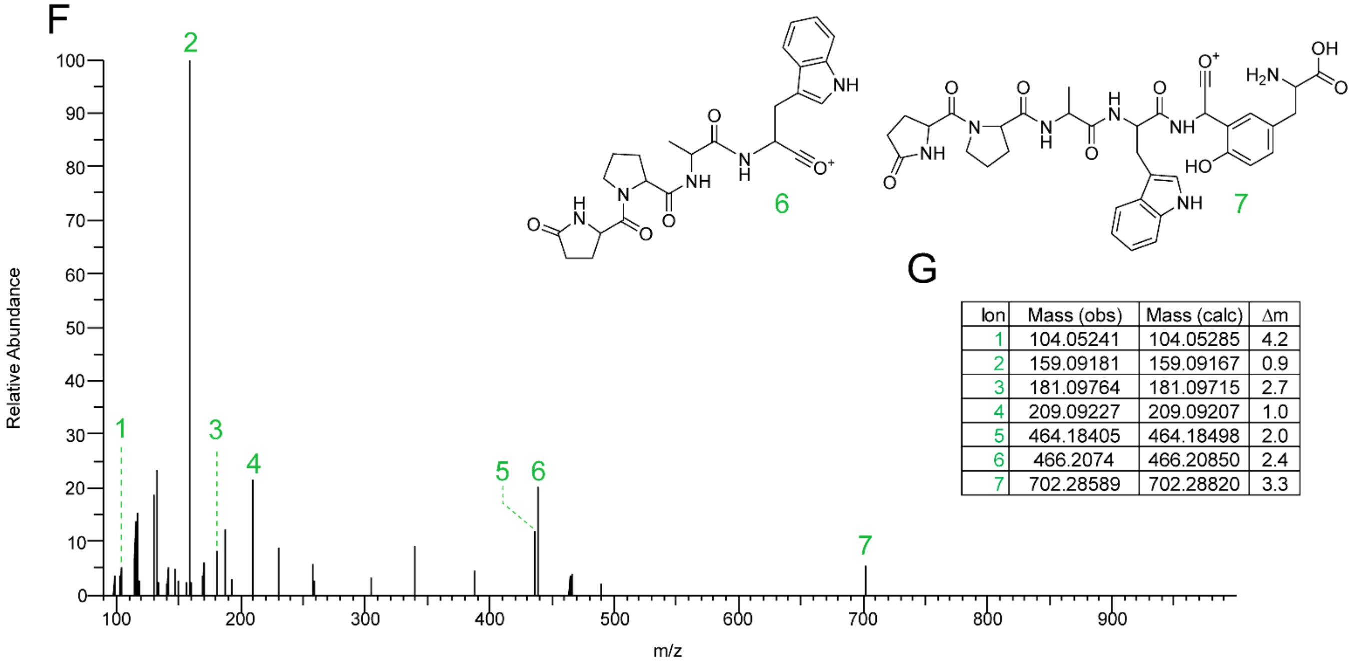


G

| Ion | Mass (obs) | Mass (calc) | Δm |
|---|---|---|---|
| 1 | 104.05241 | 104.05285 | 4.2 |
| 2 | 159.09181 | 159.09167 | 0.9 |
| 3 | 181.09764 | 181.09715 | 2.7 |
| 4 | 209.09227 | 209.09207 | 1.0 |
| 5 | 464.18405 | 464.18498 | 2.0 |
| 6 | 466.2074 | 466.20850 | 2.4 |
| 7 | 702.28589 | 702.28820 | 3.3 |

**Figure S17. MS/MS analysis of cyclo-[QPAWGLMY].** (**A**) Putative structure of cyclo-[QPAWGLMY]. (**B**) Sequence of NanaminCyclase+[QPAWGLMY]. (**C**) EIC (946.41275 m/z, 5 ppm, +p ESI Full MS, [300-1500]) of *N. benthamiana* agroinfiltrated with NanaminCyclase+[QPAWGLMY]. (**D**) MS analysis of cyclo-[QPAWGLMY] at RT 5.64 min. (**E**) Predicted daughter ions. (**F**) MS/MS spectra. (**G**) Observed vs. calculated daughter ion masses.

A

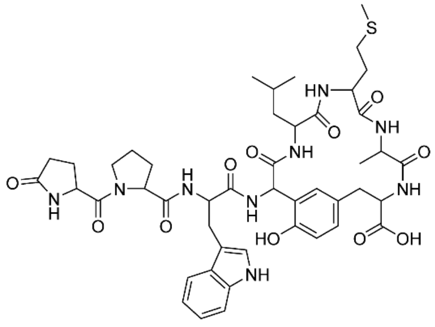


B

```
>NanaminCyclase+[QPWGLMAY]
MEFHFLLLSLAFIASHAANLSPEVYWKVKLPNTPMPKPIKDA
LTHYSAVEGEFHKLRQPWGLMAYQSATEGELHKLRQPWGLMA
YQPTTEGELHKLRQPWGLMAYQPATEGEIHKLRQPWGLMAYQ
LATEGELRKLRQPWGLMAYQGATKKDVNENDLVTPYFFETDL
HQGKKMNLFSLKNYNPAPILPRKVADSIPFSSEKIGEILNHF
SVDKDSEGAKMIEKIVKLCEEPAGTRETRYCATSLESMVDFI
SSHLGTNNILAMSTEVEKETPEAQTYIIEEVKEKTNGKGIVC
HKAAYPYAVHVCHDFGNSRTFKVSMVGADGTKVNAVSICHEN
TAAMNPKALPFQLLNIKPGGKPFCHFILDDQVAVVPAQDATQ
VAEN
```

C

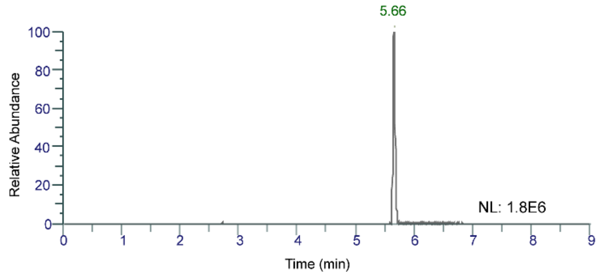


D

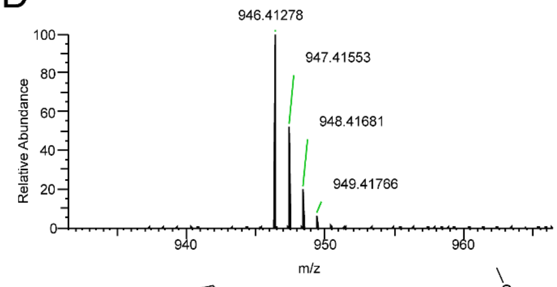


E

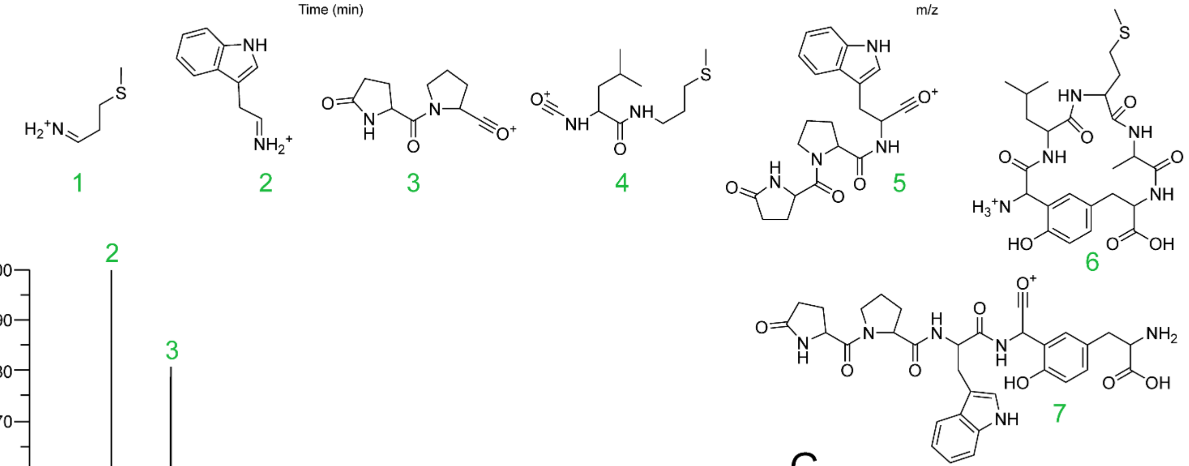


F

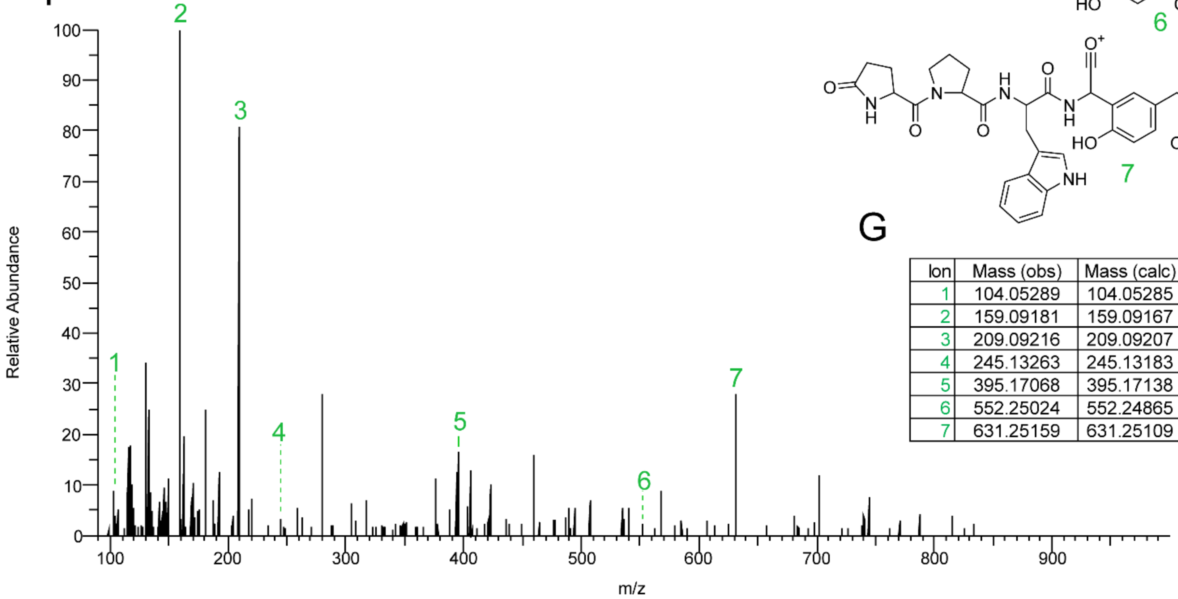


G

| Ion | Mass (obs) | Mass (calc) | Δm |
|---|---|---|---|
| 1 | 104.05289 | 104.05285 | 0.4 |
| 2 | 159.09181 | 159.09167 | 0.9 |
| 3 | 209.09216 | 209.09207 | 0.4 |
| 4 | 245.13263 | 245.13183 | 3.3 |
| 5 | 395.17068 | 395.17138 | 1.8 |
| 6 | 552.25024 | 552.24865 | 2.9 |
| 7 | 631.25159 | 631.25109 | 0.8 |

**Figure S18. MS/MS analysis of cyclo-[QPWGLMAY].** (**A**) Putative structure of cyclo-[QPWGLMAY]. (**B**) Sequence of NanaminCyclase+[QPWGLMAY]. (**C**) EIC (946.41275 m/z, 5 ppm, +p ESI Full MS, [300-1500]) of *N. benthamiana* agroinfiltrated with NanaminCyclase+[QPWGLMAY]. (**D**) MS analysis of cyclo-[QPWGLMAY] at RT 5.66 min. (**E**) Predicted daughter ions. (**F**) MS/MS spectra. (**G**) Observed vs. calculated daughter ion masses.

A

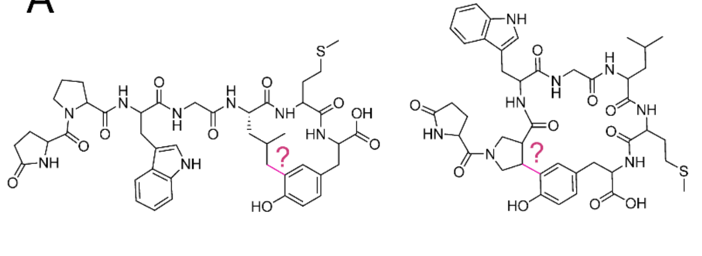


Possible cyclization sites for
Isocyclo-[QPWGLMY]

B

>LyciuminCyclase+[QPWGLMY]
MEFHFLLLSVSTLAFIASHAANLSPEVYWKVKLPNTPMPKPI
KDALTHYSEGEFHKLRQPWGLMYQSATEGELHKLRQPWGLMY
QPATKGELHKLRQPWGLMYQPATEDELHKLRQPWGLMYQPAT
EGELQKLRQPWGLMYQPATEGELHKLRVTPYFFETDLHQGKK
MNLPSLKNHNPAPILPRKIADSIPFSSDKMEEILNHFSVDKD
SEVAKMIKKTIKMCEEPAGNGEKKYCATSLESMVDFATSHLG
TNNIIALSTEVEKETPQVQIYTIEEVKEKANGKGVICHKIGY
PYAVHFCHDVGSTRTFMVSMVGADGTKVNAVAECHEDTSLMN
PKALPFQLLNVKPGDKPICHFILDDQIALVIPSRDGTQVAEN
VPAQDATQVAEN

C

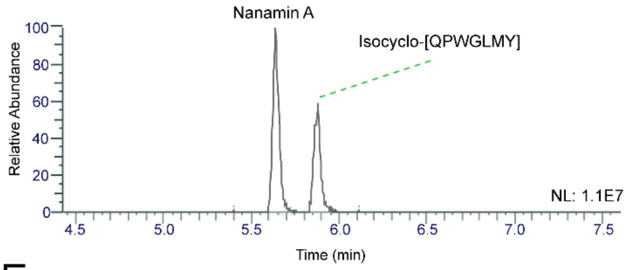


D

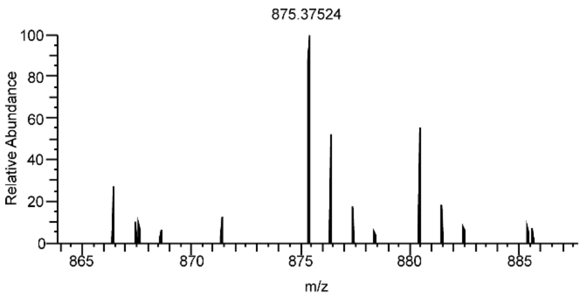


E

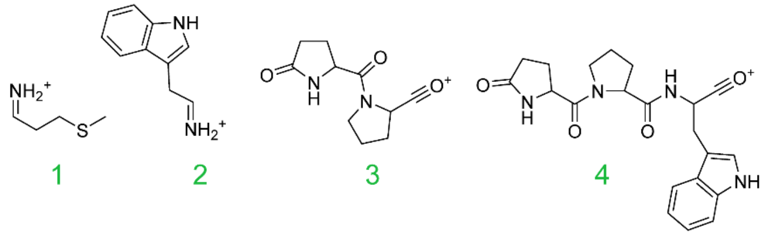


F

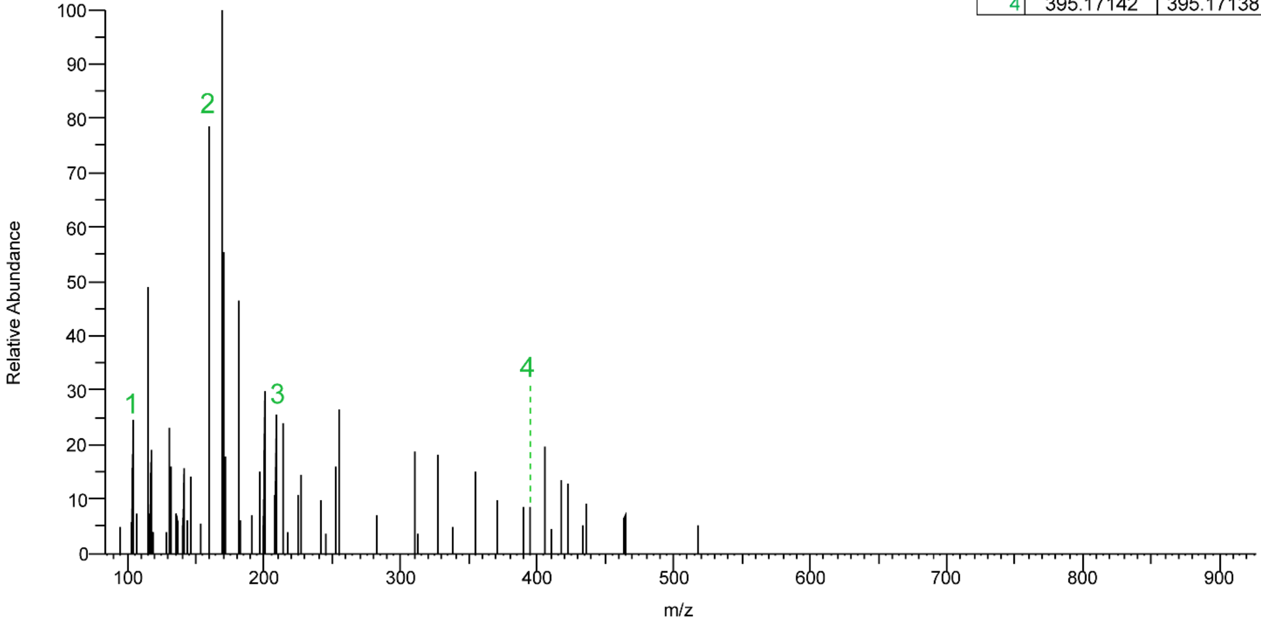


G

| Ion | Mass (obs) | Mass (calc) | Δm |
|---|---|---|---|
| 1 | 104.05280 | 104.05285 | 0.5 |
| 2 | 159.09180 | 159.09167 | 0.8 |
| 3 | 209.09204 | 209.09207 | 0.1 |
| 4 | 395.17142 | 395.17138 | 0.1 |

**Figure S19. MS/MS analysis of isocyclo-[QPWGLMY].** (**A**) Possible structures of the isocyclo-[QPWGLMY]. MS/MS analysis indicates that its cyclization is distinct from that of nanamin. (**B**) Sequence of LyciuminCyclase+[QPWGLMY]. (**C**) EIC (875.37564 m/z, 5 ppm, +p ESI Full MS, [300-1500]) of *N. benthamiana* agroinfiltrated with LyciuminCyclase+[QPWGLMY]. (**D**) MS analysis of isocyclo-[QPWGLMY] at RT 5.9 minutes. (**E**) Predicted daughter ions. (**F**) MS/MS spectra. (**G**) Observed vs. calculated daughter ion masses.

A

B

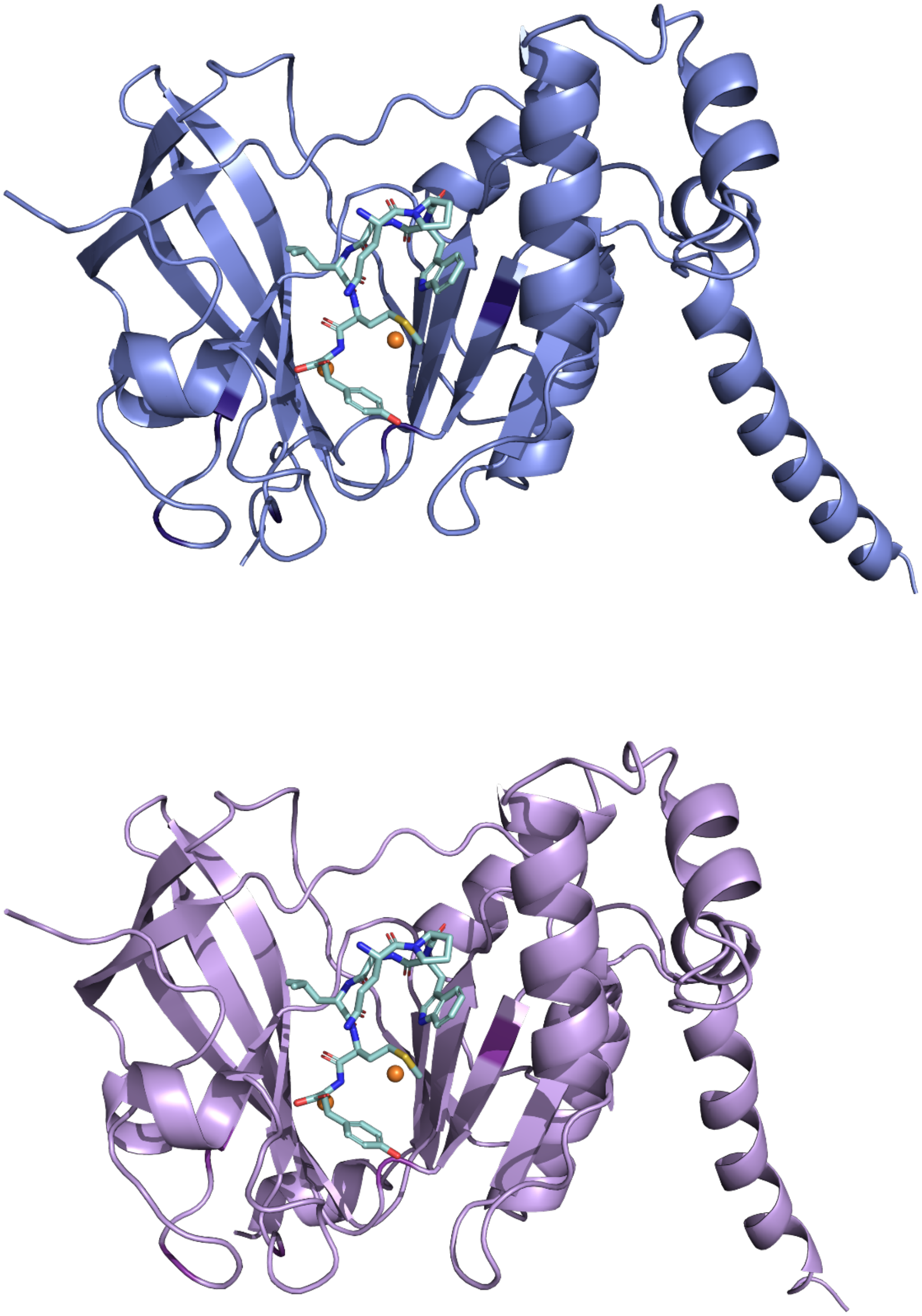

**Figure S20. Predictive AlphaFold Structures for NatBURP1 and NatBURP2.** (**A**). Predictive structure of NatBURP1, without tandem repeats. Additional inputs: two copper ions, and the peptide [QPWGLMY]. (**B**) Predictive Structure of NatBURP2, without tandem repeats. Additional inputs: two copper ions (orange), and the peptide [QPWGLMY] (teal).
